# A single cell atlas reveals distinct immune landscapes in transplant and primary tumors that determine response or resistance to immunotherapy

**DOI:** 10.1101/2020.03.11.978387

**Authors:** Amy J. Wisdom, Yvonne M. Mowery, Cierra S. Hong, Xiaodi Qin, Dadong Zhang, Jonathon E. Himes, Lan Chen, Hélène Fradin, Eric S. Muise, Eric S. Xu, David J. Carpenter, Collin L. Kent, Kimberly S. Smythe, Nerissa Williams, Lixia Luo, Yan Ma, Kouros Owzar, Todd Bradley, David G. Kirsch

**Author notes:** Corresponding Authors (YMM, DGK).

## Abstract

Despite impressive responses in some patients, immunotherapy fails to cure most cancer patients. Preclinical studies indicate that radiotherapy synergizes with immunotherapy, promoting radiation-induced antitumor immunity. Nearly all preclinical immunotherapy studies utilize transplant tumor models, but cure rates of transplant tumor models treated with immunotherapy often overestimate patient responses. Here, we show that transplant tumors are cured by PD-1 blockade and radiotherapy, but identical treatment fails in autochthonous tumors. We generated a single-cell atlas of tumor-infiltrating immune cells from transplant and primary tumors treated with radiation and immunotherapy, which reveals striking differences in their immune landscapes. Although radiotherapy remodels myeloid cell phenotypes in primary and transplant tumors, only transplant tumors are enriched for CD8+ T cells that mediate tumor clearance while mice with primary sarcomas demonstrate tumor-specific tolerance. These results identify distinct microenvironments in tumors that coevolve with the immune system, which promote tolerance that must be overcome for immune-mediated cancer cures.

---

Many cancer patients receive radiation therapy (RT) for palliation or with the intent to achieve permanent regression of the irradiated tumor^1^. Preclinical studies using transplanted tumor models demonstrate that focal RT can synergize with immune checkpoint inhibitors to generate systemic immune responses. In these abscopal responses, RT acts as an *in situ* vaccine^2–4^ to eliminate tumors outside of the radiation field in a T cell- and type I interferon-dependent manner^5–7^. Preclinical studies with transplanted tumors demonstrating high cure rates with checkpoint blockade and radiotherapy^8–11^ have led to numerous clinical trials^12, 13^, but emerging results are disappointing^9, 14–16^.

To gain insight into mechanisms of response and resistance to immune checkpoint blockade and radiotherapy, we administered RT and anti-programmed cell death-1 (PD-1) antibody to mice bearing primary or transplant tumors from a novel high-mutation mouse model of sarcoma^17^. Like other studies with transplanted tumors that do not develop in their native microenvironment under immunosurveillance, we find that transplanted tumors in syngeneic mice are cured by immune checkpoint blockade and RT. However, the identical treatment fails to achieve local control in autochthonous sarcomas from the same model system.

Using single-cell RNA sequencing and mass cytometric profiling, we generated a single cell atlas of tumor-infiltrating immune cells from transplant and primary tumors before and after radiation and anti-PD-1 immunotherapy, which reveals marked differences in their immune landscapes. Furthermore, we show that mice that previously developed a primary tumor demonstrate immune tolerance to their own tumors after auto-transplantation, but reject transplanted tumors from other mice. Because auto-transplantation fails to generate a response to anti-PD-1 immunotherapy, our observations support a model in which coevolution of tumors and the immune system generates an immune cell landscape that favors tumor tolerance. Furthermore, we identify features of immune resistance that exist specifically in treatment-resistant primary tumors that must be targeted for radiation and immunotherapy to realize their full therapeutic potential.

## Results

### Responses to radiation and PD-1 blockade

Because human cancers with a higher tumor mutational burden may be more likely to respond to immune checkpoint inhibition^18–20^, we tested the efficacy of RT and an antibody targeting PD-1 in a high mutational load mouse model of sarcoma^17^. We injected the gastrocnemius muscle of *Trp53^fl/fl^* mice with an adenovirus expressing Cre recombinase (Adeno-Cre) to delete *Trp53*, followed by injection with the carcinogen 3-methylcholanthrene (MCA). Primary p53/MCA sarcomas developed at the injection site under the selective pressure of immunoediting in immunocompetent mice^17^. A cell line from an untreated primary p53/MCA sarcoma was transplanted into the gastrocnemius muscle of syngeneic mice. The resulting tumors were cured by PD-1 blockade and 20 Gy RT (**Fig. 1a**). However, the same combination treatment failed to cure primary p53/MCA sarcomas (**Fig. 1b**), even with the addition of CTLA-4 blockade (**Supplementary Fig. 1a, b**). Transplant tumor cure by anti-PD-1 and RT was both CD4- and CD8-dependent (**Supplementary Fig. 1c, d**).

**Fig. 1.**
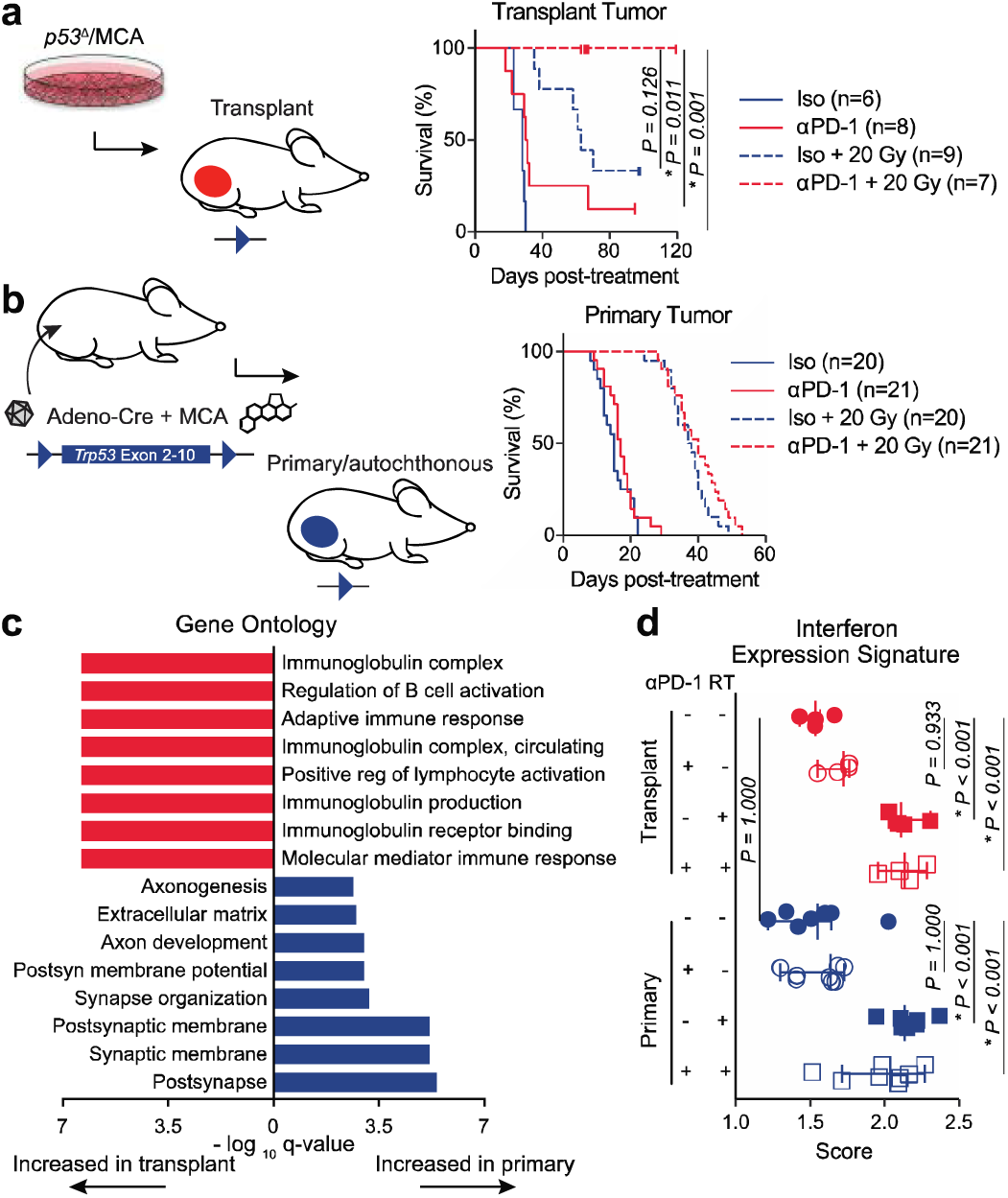
Immune checkpoint blockade and radiation therapy cure transplant but not primary sarcomas. **a**, Transplant tumor initiation by p53/MCA cell injection into the gastrocnemius. Mice were treated with anti (α)-PD-1 (red) or isotype control (blue) antibody and 0 (solid) or 20 (dashed) Gy. **b**, Primary sarcoma initiation by intramuscular injection of Adeno-Cre and MCA. Treatment as in panel **a**. Survival curves estimated using Kaplan-Meier method; pairwise significance determined by log-rank test and Bonferroni correction. **c**, Gene ontology analysis of differentially regulated processes in primary (blue) versus transplant (red) sarcomas after isotype treatment. **d**, Expression of interferon stimulated gene expression signature after treatment with αPD-1 or RT (20 Gy) in primary (blue) and transplant (red) tumors. n = 4 per group for transplant tumors, n = 8 per group for primary tumors. Significance determined by Wilcoxon rank-sum test and Bonferroni correction.

To assess transcriptional differences in primary and transplant sarcomas, we performed RNA sequencing on bulk tumor RNA. Primary and transplant sarcomas were harvested 3 days after treatment with either 0 or 20 Gy and anti-PD-1 or isotype control antibody. Notably, principal components analysis showed that tumor type (primary vs. transplant) was the major factor driving transcriptional differences, rather than treatment with radiation or anti-PD-1 therapy (**Supplementary Fig. 1e**). Comparing the gene expression differences in transplant and primary sarcomas revealed that, within the many differentially expressed genes (**Supplementary Fig. 1f, g**), primary tumors expressed more axonal and developmental gene sets, while transplant tumors exhibited enrichment of immune-related genes even at baseline (Isotype + 0 Gy) (**Fig. 1c**). Expression of an interferon-stimulated gene (ISG) signature^21^ was similar between primary and transplant tumors (**Fig. 1d**), suggesting that the level of inflammation in the tumor microenvironment did not differ significantly before treatment. Additionally, expression of the ISG signature after radiation therapy with or without anti-PD-1 therapy increased by a similar magnitude in both tumor models. These findings suggest that specific populations of the immune system mediate the differences in treatment response to anti-PD-1 and RT.

### Immune responses to anti-PD-1 therapy

Tumor-infiltrating CD8+ T cells correlate with the likelihood of response to anti-PD-1 therapy and increase during anti-PD-1 treatment in responsive tumors^22, 23^. Therefore, we examined CD8+ T cells by immunohistochemistry in primary and transplant tumors three days after treatment with anti-PD-1 antibody, which revealed similar numbers of CD8+ T cells at baseline but increased CD8+ T cells after anti-PD-1 therapy in transplant tumors (**Supplementary Fig. 2a, b**). To further study the temporal dynamics of CD8+ T cells and other immune cell populations in primary and transplant tumors, we used a panel of 37 heavy metal-conjugated antibodies to analyze independent tumor samples by mass cytometry (CyTOF)^24^ one, three, and seven days after treatment with anti-PD-1, RT, or anti-PD-1 and RT (**Supplementary Fig. 2c-g**). We again found similar numbers of CD8+ T cells in primary tumors at baseline (**Supplementary Fig. 2c**), suggesting that T cell exclusion was not a key mechanism for tumor resistance. However, CyTOF profiling revealed a subset of activated CD8+ T cells expressing Granzyme B (GzmB), Ki67, and the immune checkpoints Lag3 and Tim3 that was significantly enriched in transplant tumors (**Supplementary Fig. 2g**). These results suggest that primary and transplant tumors promote distinct phenotypes of CD8+ T cells.

To gain insight into the transcriptional differences in CD8+ T cells in primary and transplant tumors, we performed single-cell RNA sequencing (scRNA-seq) on FACS-sorted CD45+ tumor-infiltrating immune cells from sarcomas harvested 3 days after treatment with either anti-PD-1 antibody or isotype control (primary and transplant) and 0 or 20 Gy (primary tumors only). After filtering and quality control, scRNA-seq analysis yielded data for 98,219 cells with 52,220 mean reads per cell detecting 1,570 median genes per cell. We performed tSNE dimensionality reduction and graph-based clustering of all cells in aggregate to identify cells with distinct transcriptional profiles (**Fig. 2a**). Graph-based clustering identified 23 cell clusters with distinct transcriptional programs that could be readily assigned to known cell lineages using marker genes (**Fig. 2a**), resulting in classification of two B cell clusters (C19, C22), nine myeloid cell clusters (C0, C1, C2, C3, C4, C6, C8, C11, and C18), six T/NK cell clusters (C5, C7, C9, C12, C13, C16), two clusters of conventional dendritic cells (cDC) (C20, C21), one cluster of plasmacytoid dendritic cells (pDC) (C14), one cluster of neutrophils (C10), and one cluster of fibroblasts (C17), which was subsequently eliminated from the analysis.

**Fig. 2.**
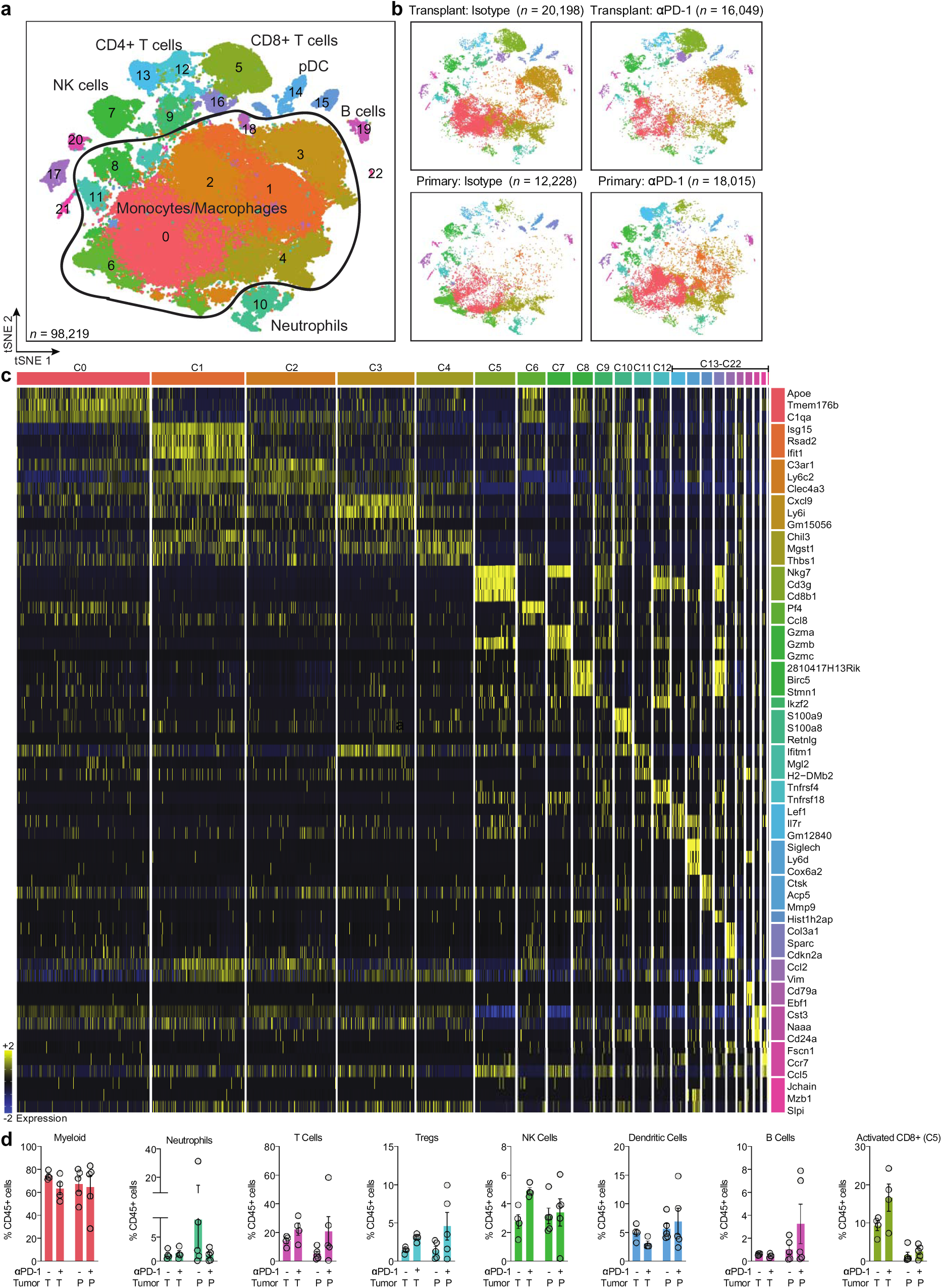
Characterization of immune microenvironment by single-cell RNA sequencing. **a**, CD45+ immune cells from all tumor and treatment groups, overlaid on tSNE representation and color coded by cluster. Cell populations expressing genes consistent with monocyte/macrophage phenotype are outlined. **b**, Distribution of immune cells separated by indicated tumor and treatment type**. c**, Scaled expression values of top three discriminative genes per cluster. **d**, Bar graphs comparing frequency of indicated cell type (% CD45+) from αPD-1 (+) or isotype control (-). Each symbol represents an individual mouse; n = 4 per group for transplant tumors, n = 5 per group for primary tumors. Data show mean ± SEM.

The immune cell atlases generated by the complementary methods of CyTOF and scRNA-seq revealed significant differences between primary and transplant tumors samples in all major immune cell populations (**Fig. 2b-d, Supplementary Fig. 2c, d**). In fact, multiple cell populations were enriched specifically in primary or transplant sarcomas. While transplant tumors demonstrated robust infiltration of activated CD8+ T cells (C5) by scRNA-seq, this CD8+ lymphocyte population was nearly absent in primary tumors, even after anti-PD-1 therapy (**Fig. 2b, d**). Different myeloid cell clusters were also enriched specifically in primary or transplant tumors (**Fig 2b**), reflecting the transcriptional differences within the myeloid cell compartments of these models. Using these complementary approaches, we found marked differences by tumor type in tumor-infiltrating immune cells based on transcriptional profile, population distribution, and protein expression, which were accentuated by remodeling after anti-PD-1 therapy.

### Lymphocyte remodeling in primary tumors

To increase the resolution and more accurately define the lymphoid populations identified by scRNA-seq, we computationally separated 14,705 lymphoid cells from all CD45+ cells for further analysis (**Fig. 3a**). This approach yielded 12 distinct lymphoid subpopulations, including regulatory T cells (Treg) (L5), naive CD4+ T cells (L9), CD8+ T cells (L0, L4, L6, and L8), one population containing both CD4+ and CD8+ memory T cells (L3), natural killer cells (L1, L10), B cells (L8), and plasma cells (L12) (**Fig. 3a-c** and **Supplementary Fig. 3a, b, 4a**).

**Fig. 3.**
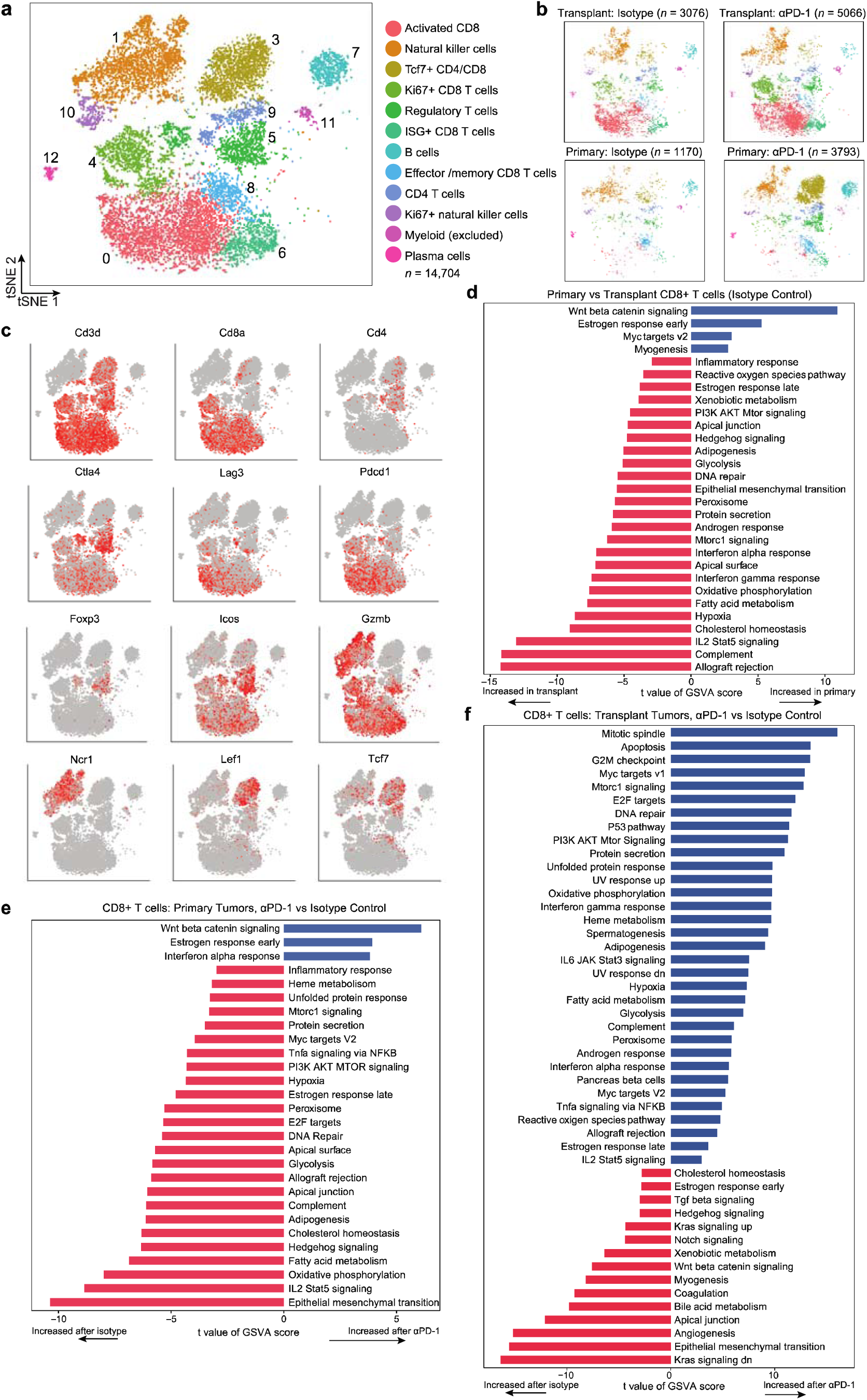
Remodeling of tumor-infiltrating lymphoid populations. **a**, tSNE plot of lymphoid cell scRNA-seq subclustering from all tumor and treatment groups. **b**, Distribution of immune cells, separated by tumor and treatment type. n = 4 tumors per group for transplant, n = 5 tumors per group for primary. **c**, Expression levels of marker genes. **d**, Pathways with significantly different activities per cell using gene set variation analysis (GSVA) between CD8+ T cells from primary vs transplant tumors treated with isotype control antibody (n = 2,572 cells analyzed). **e**, Pathways with significantly different activities per cell using GSVA between CD8+ T cells from primary tumors treated with αPD-1 vs isotype control antibody (n = 2,473 cells analyzed). **f**, Pathways with significantly different activities per cell using GSVA between CD8+ T cells from transplant αPD-1 vs isotype control antibody (n = 5,828 cells analyzed). For **d-f**, Pathways shown are significant at false discovery rate < 0.01.

The most apparent difference was the low number of activated CD8+ T cells (L0) in primary tumors relative to transplant tumors, even after treatment with anti-PD-1 therapy (**Fig. 3b and Supplementary Figure 3b**). Within the CD8+ T cell populations, L0, L4, and L6 were enriched in transplant tumors, while population L8 was more abundant in primary tumors (**Fig. 3b and Supplementary Figure 3b**). The majority of CD8+ T cells from transplant tumors fell into population L0, which expressed high levels of genes associated with T cell activation and/or exhaustion (*Pdcd1, Havcr2, Lag3, Ctla4, Cd38, Entpd1)* (**Fig. 3c, d, Supplementary 3b, 4b, and Table S1**). Actively cycling CD8+ T cells (*Cdk1, Ccnb1, Ccna2, Mki67, Cdk4*) within population L4 were also more abundant within transplant tumors and, consistent with previous studies^25^, increased after anti-PD-1 treatment (**Fig. 3b, Supplementary Fig. 3b, and Table S1**). Population L6 expressed high levels of genes associated with the type I interferon response (*Stat1, Irf7*, *Mx1, Ifit1*, *Isg15*, *Ccl5*) and moderate levels of the same exhaustion markers found in L0 (**Supplementary Fig. 4b and Table S1**). L8, a memory population which lacked exhaustion markers and primarily expressed markers associated with survival/proliferation (*Il7r, Slamf6, Ccnd3, Ccnd2, Cd69*) was the only CD8+ population more abundant in primary tumors than transplant tumors (**Supplementary Fig. 3b, 4b, and Table S1**).

In chronic viral infection and cancer, a subset of exhausted T cells expressing high levels of PD-1 and other inhibitory checkpoints can proliferate and differentiate into effectors that mediate long-term immune control after anti-PD-1 immunotherapy^26, 27^. In the absence of treatment, CD8+ T cells from transplant tumors exhibited significantly different gene expression profiles compared to those in primary tumors (**Fig. 3d**) and resembled the previously described exhausted T cell population. CD8+ T cells in transplant tumors exhibited higher levels of activation/exhaustion markers (*Gzmb, Lag3, Havcr2, Tnfrsf9, Icos, Pdcd1*) and increased activity of transplant rejection and interferon response pathways (**Fig. 3c, d, and Table S2**). Comparison of pathway activity in CD8+ T cells revealed elevated Wnt/β-Catenin and Myc target gene signaling in primary tumors compared to transplant tumors at baseline (isotype control treatment) (**Fig. 3d**). Interestingly, the activity of the Wnt/β-Catenin signaling pathway in CD8+ T cells increased further after anti-PD-1 treatment in primary tumors (**Fig. 3e**) but decreased after anti-PD-1 treatment in transplant tumors (**Fig. 3f**). Lymphocyte overexpression of Wnt/β-Catenin target genes has been shown to lead to apoptosis in mature T cells^28^, and tumor cell-intrinsic β-Catenin signaling is associated with T cell exclusion and immune evasion^29^.

Within CD8+ T cells in transplant tumors, treatment with anti-PD-1 antibody induced higher levels of granzyme expression (*Gzma, Gzmb*) and cell proliferation genes (*Rps12, Rpl5, Eif4a1, Top2a*) and reduced expression of the exhaustion markers *Tnfrsf18* (*Gitr*) and *Lag3*. Within CD8+ T cells in primary tumors, anti-PD-1 therapy had similar effects on granzyme expression (increased *Gzma, Gzmb*), proliferation genes (increased *Rps12*, *Rpl5, and Smchd1*), and exhaustion markers (decreased *Tnfrsf18* and *Pdcd1*) (**Supplementary Fig. 4b and Table S2**). Interestingly, in the small fraction of CD8+ T cells from primary tumors in L0, treatment with anti-PD-1 antibody induced high expression of *Tox* (**Supplementary Fig. 4b and Table S2**), a critical regulator of tumor-specific T cell differentiation that promotes T cell commitment to an exhausted phenotype^30–32^. These analyses suggest that the anti-PD-1 antibody was engaging T cells in both models, but specific transcriptional CD8+ T cell states are associated with primary tumor resistance to anti-PD-1 immunotherapy.

## Myeloid remodeling by anti-PD-1 therapy

Tumor-infiltrating macrophages can promote tumor progression through surface expression of immune checkpoints and by production of anti-inflammatory cytokines that induce immune suppression and resistance to checkpoint inhibition. In transplant tumor models, successful blockade of the PD-1/PD-L1 pathway can repolarize macrophages toward a proinflammatory phenotype characterized by a decrease in M2 markers such as Arginase-1^33, 34^. Tumor-infiltrating myeloid cells comprise the largest fraction of immune cells in both primary and transplant tumors, and they undergo significant remodeling with anti-PD-1 therapy (**Fig. 2b, Supplementary Fig. 5** and **6**). Using CyTOF, we found that PD-L1+ macrophages were more abundant in early-stage transplant tumors than in primary tumors (**Fig. 4a**), suggesting a possible mechanism for transplant tumor response to anti-PD-1 therapy. To interrogate additional transcriptional differences in the myeloid phenotypes of primary and transplant sarcomas, we sub-clustered the 73,039 myeloid cells identified by scRNA-seq. This analysis (**Fig. 4b, c, and Supplementary Fig. 5**) yielded 14 myeloid subpopulations, which we compared to published data sets for cell type identification (**Supplementary Fig. 6a**). Enrichment of myeloid cell clusters specifically in primary or transplant tumors (**Fig. 4c**) reflects the transcriptional differences within the myeloid cell compartments of these models, while the incomplete separation between clusters suggests overlapping and highly plastic myeloid cell phenotypes.

**Fig. 4.**
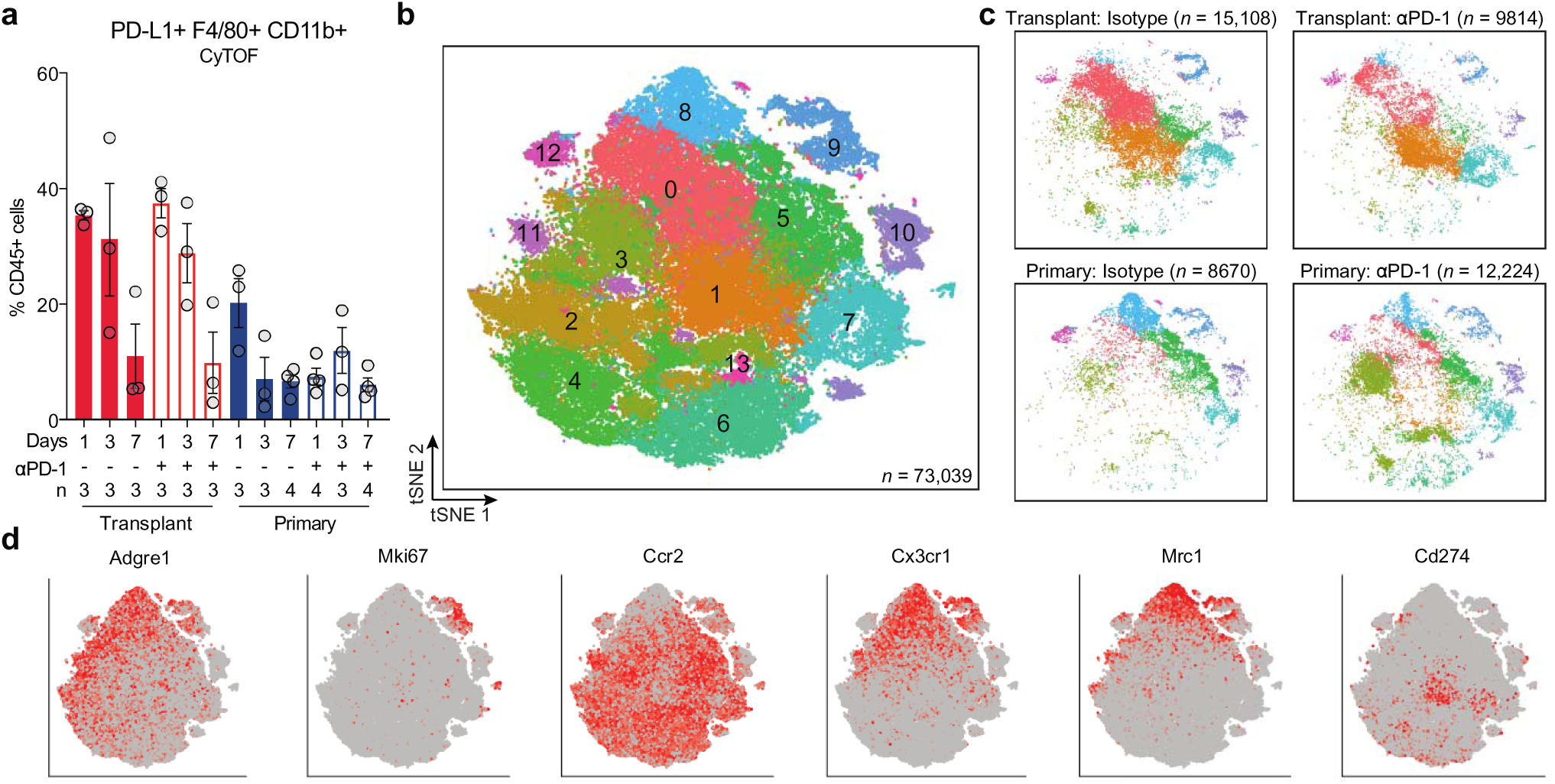
Defining tumor-infiltrating myeloid cells. **a**, Frequency of PD-L1+ F4/80+ CD11b+ macrophages by CyTOF. **b**, tSNE plot of myeloid cell scRNA-seq subclustering from all tumor and treatment groups. **c**, Distribution of myeloid cells, separated by tumor and treatment type. **d**, Expression levels of marker genes.

The majority of myeloid cells from untreated (isotype) transplant tumors fell into clusters Y0, Y1, Y5, and Y7 (**Supplementary Fig. 5b**). Cluster Y0 expressed high levels of genes related to complement (*C1qc, C1qa, C1qb*) and interferon-stimulated cytokines (*Cxcl14, Cxcl16, Ccl3, Ccl4*). Cells in Y1 expressed classical macrophage markers, including *Itga4* and *Il1r2* (**Supplementary Fig. 5b and Table S1**). Additionally, Y1 expressed high levels of proinflammatory transcription factors (*Stat1* and *Irf1)* that cooperate to increase transcription of *Cd274* (PD-L1)^35^, which was also highly expressed in this cluster. Additional populations of cells from untreated transplant tumors clustered into Y5 and Y7, which expressed immunosuppressive genes, including *Ptgs2*, *Ccr2*, *Chil3*, and *Tgfb*, consistent with M2 macrophage phenotypes (**Supplementary Fig. 5b, 6b, Table S1**).

Upon treatment with anti-PD1 therapy, the frequency of transplant-infiltrating myeloid cells in Y0 decreased, which was accompanied by an increase in both proinflammatory macrophages (Y1, representing 42% of all myeloid cells) and anti-inflammatory macrophages (Y7, 16%) (**Supplementary Fig. 5b**). Anti-PD-1 treatment led to a further increase in *Stat1*, *Irf1*, and *Cd274*, which was accompanied by transcriptional downregulation of the anti-inflammatory macrophage marker *Cx3cr1* (**Supplementary Fig. 6b and Table S3**). These findings support a model where the transcriptional response to anti-PD-1 therapy in transplant-infiltrating myeloid cells is dominated by interferon gamma^25^.

Within primary tumors treated with isotype control antibody, the majority of myeloid cells clustered into Y8, Y5, Y7, and Y3 (**Fig. 4c**). Y8 macrophages expressed *Csf1r* and *Adgre1,* and they had the highest levels of the anti-inflammatory genes *Mrc1, Cx3cr1,* and *Mertk* (**Supplementary Fig. 6b**). Y5, which represented the majority of myeloid cells in progressively growing primary tumors, expressed *Ccr2* and *Ly6c2*, consistent with a monocyte phenotype, as well as high levels of *Ptgs2*. Y8, Y5, and Y7 macrophages, which all represent immunosuppressive myeloid populations, represented the majority of myeloid cells (> 60%) in untreated primary tumors (**Fig. 4c**). Proinflammatory myeloid cells in primary tumors fell into cluster Y3, which expressed genes associated with antigen presentation and classical macrophage activation (*Cd74, Ccr2, H2-Aa, Cd72*) but lacked the *Stat1*/*Irf1* gene expression signature seen in the proinflammatory myeloid cells from transplants in cluster Y1 (**Supplementary Fig. 6b and Table S3**). However, Y3 myeloid cells in primary tumors did express high levels of *Irf7*, a crucial regulator of the type I interferon response and monocyte-to-macrophage differentiation^36^.

Treatment of primary tumors with anti-PD-1 therapy decreased the frequency of immunosuppressive myeloid cells in Y5 and Y8 and led to an increase in proinflammatory macrophages (Y3) (**Fig. 4c**). Within myeloid cells in primary tumors, PD-1 blockade led to significantly decreased expression of *Mrc1* and *Ptgs2*, as well as increased expression of genes encoding antigen processing machinery (*Tap1, Tapbp, B2m*) and genes involved in the type I interferon response (*Irf7, Isg15, Ifit1, Ifit3, Ccl5*) (**Supplementary Fig. 6b and Table S3**). PD-1 blockade also induced *Stat1* and *Irf1*, suggesting that despite the immunosuppressive myeloid cell environment in isotype control-treated primary tumors, treatment with PD-1 blockade successfully induces myeloid cell phenotype molding toward an anti-tumor phenotype.

### Immune responses to radiation therapy

In preclinical studies using transplanted tumor models, focal RT can synergize with immune checkpoint inhibitors by increasing tumor immunogenicity and by reinvigorating the anti-tumor immune response^9, 12, 37^. To examine the immunologic responses to RT alone or combined with PD-1 blockade, we performed CyTOF on primary and transplant sarcomas 3 days after treatment with 20 Gy focal tumor RT and either anti-PD-1 or isotype control antibody. RT induced extensive remodeling across both lymphoid and myeloid cell compartments (**Supplementary Fig. 7a**). Specifically, CyTOF revealed that RT significantly reduced immunosuppressive M2 macrophages (F4/80+ CD206+ CX3CR1+) at early time points in both primary and transplant sarcomas, with and without anti-PD-1 therapy (**Fig. 5a, Supplementary Fig. 7b, c**).

**Fig. 5.**
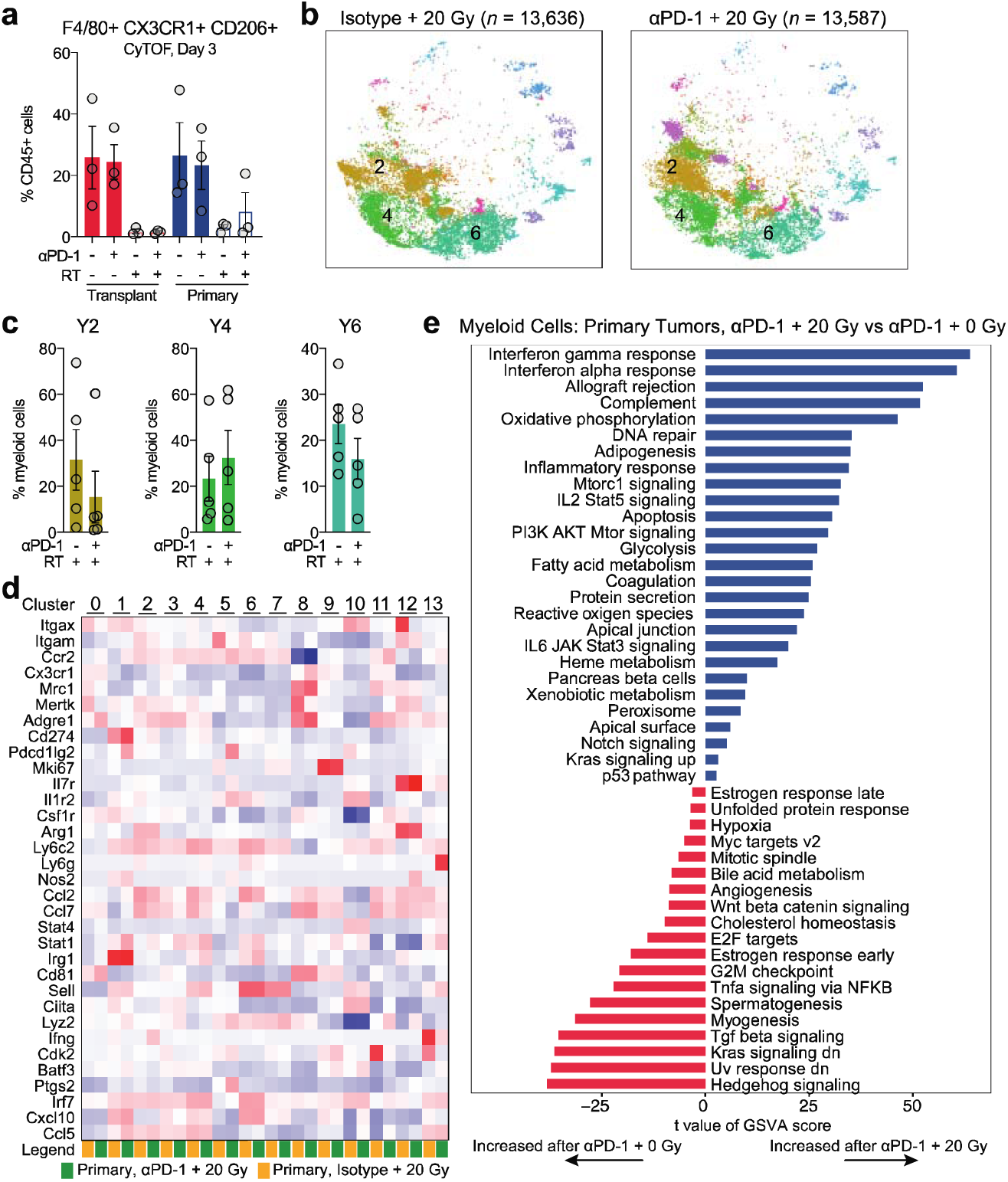
Immunologic responses to combination PD-1 blockade and radiotherapy. **a**, Frequency of CX3CR1+ CD206+ macrophages by CyTOF. **b**, Distribution of myeloid cells by scRNA-seq for primary tumors after indicated treatment. **c**, Frequency of indicated cluster population (% myeloid cells) after 20 Gy radiation therapy and either isotype control or αPD-1 antibody, analyzed by scRNA-seq. Each symbol represents an individual mouse. n = 4 per group for transplant tumors, n = 5 per group for primary tumors. Data show mean ± SEM. **d**, Heat map comparing expression of genes in indicated clusters and treatment groups. **e**, Pathways with significantly different activities per cell using GSVA between myeloid cells from primary tumors treated αPD-1 and 20 Gy vs 0 Gy radiation (n = 25,811 cells analyzed). Pathways shown are significant at false discovery rate < 0.01.

**Fig. 6.**
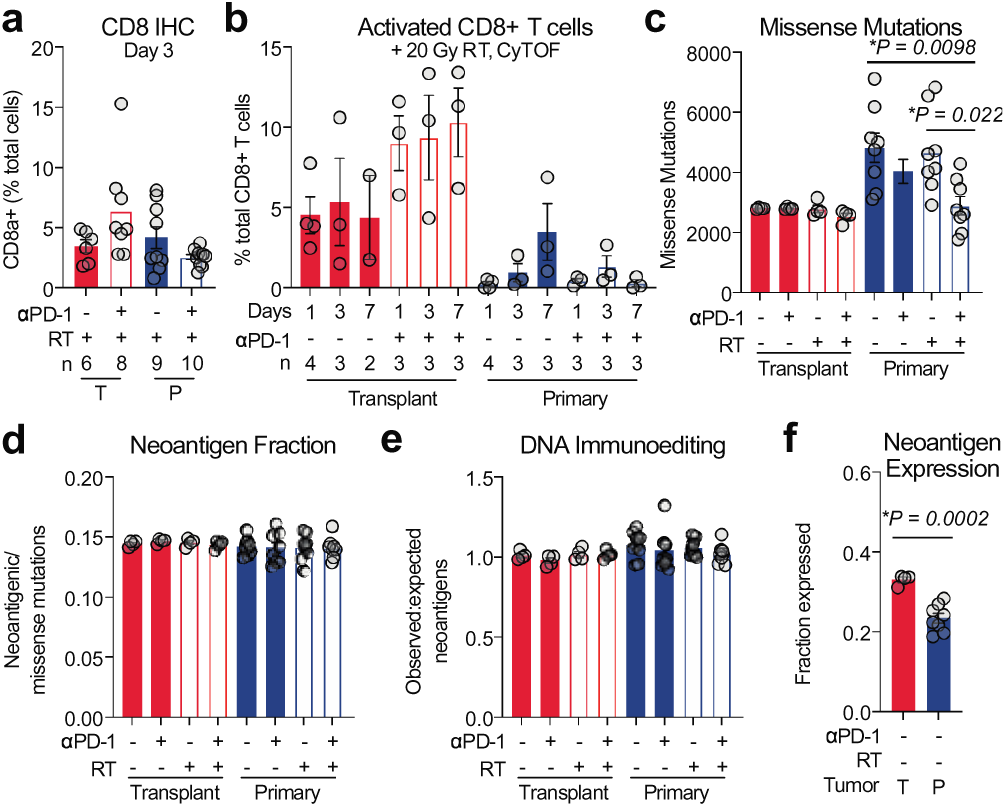
Lymphocyte and genomic responses to PD-1 blockade and radiotherapy. **a**, Percent CD8a+ cells by immunohistochemistry 3 days after treatment. T, transplant; P, primary. **b**, Frequency of activated (Tim3+ Lag3+ Ki67+ GzmB+) CD8+ T cells by CyTOF. **c**, Missense mutations in transplant and primary tumors harvested 3 days after specified treatment. **d**, Fraction of predicted neoantigenic mutations in tumors by WES. **e**, Ratio of observed:expected neoantigens. **f**, Fraction of neoantigenic mutations expressed (> 5 reads) by RNA sequencing. Each symbol represents an individual mouse, n = 4 per group for transplant tumors, n = 8 per group for primary tumors. Data show mean ± SEM. Significance determined by one-way ANOVA with Tukey’s multiple comparisons test in **c** and two-sided Student’s t-test in **f**.

To examine the transcriptional effects of RT on the immune microenvironment of radiation-resistant primary tumors, we performed scRNA-seq on CD45+ cells isolated from primary sarcomas 3 days after treatment with RT and either anti-PD-1 or isotype control antibody. Compared to unirradiated primary sarcomas (**Fig. 2b**), RT reshaped the transcriptome and distribution of multiple cell types, most notably the myeloid cell clusters (**Fig. 5b and Supplementary Fig. 7d**). After RT, most myeloid cells from tumors treated with isotype control antibody clustered into myeloid subclusters Y2, Y4, and Y6 (**Fig. 5c**). Y2 macrophages expressed high levels of *Ccl2* and *Ly6C* and expressed genes consistent with active phagocytosis and antibody-dependent cell-mediated cytotoxicity (*Fcgr2b, Fcgr1, Fcgr3*) and antigen/protein processing (*B2m, Ctsl, Ctsd, Ctsb, Ctss*) (**Table S1**). Y2 macrophages also expressed *Mertk*, a tyrosine kinase that can be activated by RT to cause suppressive differentiation of macrophages^38^. Clusters Y4 and Y6 expressed high levels of *Stat1, Stat2, Irf5,* and *Irf7*, likely responsible for the increased expression of interferon-related genes (*Mx1, Ifit1, Ifit3, Ifit3b, Ifit2*) and genes involved in antigen processing and presentation (*Tapbp, Tap1, Scimp*) (**Fig. 5d and Table S1**). Y4 could be distinguished from Y6 by elevated expression of *Ccl5*, while Y6 had elevated expression of the M1-associated gene *Cxcl10* (**Fig. 5d and Table S1**).

The addition of anti-PD-1 therapy to 20 Gy caused a modest decrease in cells clustering into Y2 and Y6 and a small increase in Y4 (**Fig. 5b, c**). Compared to anti-PD-1 antibody alone, RT increased activity of interferon response pathways and decreased activity of TGF-β signaling, which has been associated with resistance to immune checkpoint blockade^39^ (**Fig. 5e**).

Because RT and anti-PD-1 therapy decreased immunosuppressive macrophages (**Fig. 5a**) and upregulated proinflammatory transcripts (**Fig. 5d, e**) in primary tumor-infiltrating myeloid cells, we examined T cell responses after RT, with and without anti-PD-1 treatment. Interestingly, immunohistochemistry revealed that RT abrogated the PD-1 blockade-induced increase in CD8+ T cells in transplant tumors 3 days after treatment (**Fig. 6a and Supplementary Fig. 2a**). Using CyTOF, we found that the activated CD8+ T cells enriched in transplant tumors (**Supplementary Fig. 2g**) were slightly reduced at early time points after RT, but increased with the addition of anti-PD-1 treatment (**Fig. 6b**). However, despite the immunostimulatory effects of RT and PD-1 blockade on the immune microenvironment of primary tumors, activated CD8+ T cells did not accumulate in primary tumors after treatment (**Fig 6b**).

## Tumor-intrinsic immune evasion

Resistance to immunotherapy can be caused by both an immunosuppressive microenvironment and tumor cell-intrinsic immune evasion mechanisms^40, 41^. We performed whole exome sequencing (WES) and RNA-seq on tumor samples to identify possible genomic and transcriptomic mechanisms of tumor cell-mediated immune evasion. In primary and transplant tumors harvested 3 days after treatment with anti-PD-1 or isotype control antibody and 0 or 20 Gy, we compared paired WES data from the tumor and liver of each mouse to identify somatic mutations within each tumor.

In primary tumors, treatment with anti-PD-1 antibody decreased the number of missense mutations by approximately 15%, and the addition of 20 Gy resulted in approximately a 40% decrease in missense mutations (**Fig. 6c**). To examine whether there was evidence for immune evasion in primary tumors, we computationally examined tumor neoantigens. Despite the difference in mutation number (**Fig. 6c**), the detected frequency of neoantigenic mutations was similar across all treatment groups (**Fig. 6d**). DNA immunoediting was also similar between primary and transplant tumors (**Fig. 6e**). However, a significantly lower fraction of neoantigenic mutations was expressed within untreated (isotype + 0 Gy) primary tumors compared to transplant tumors (**Fig. 6f**). Transcriptional repression of tumor neoantigens, which occurs in human cancers^41^, is an important mechanism for immune escape in tumors that coevolve with the immune system and may not be fully recapitulated by transplant tumor models.

## Primary tumors drive immune tolerance

To determine whether the process of cell line transplantation, which may lead to elevated transcription of neoantigens, sensitizes tumors to RT and immunotherapy, we performed a series of complementary transplantation experiments (**Fig. 7**). First, we generated primary p53/MCA sarcomas in mice and amputated the tumor-bearing limb when the tumor reached ∼70 mm^3^. We then generated a cell line from each amputated tumor and approximately 1 month later, transplanted this cell line into the intact hind limb of the mouse from which the cell line was derived (i.e. donor mouse), as well as into naive syngeneic mice (**Fig. 7a**). Tumors grew out with 100% penetrance and significantly decreased latency when transplanted into the donor mice from which the tumor cell lines were derived, compared to transplantation into naive mice (**Fig. 7b**). Transplant tumors in donor mice were resistant to tumor cure by anti-PD-1 and RT. When the same tumor cell lines were injected into naive mice and treated with anti-PD-1 and RT, more than half of the mice (52%) were cured (**Fig. 7c**). In contrast to tumor cell lines derived from the same mouse (“self”), these non-self tumor cell lines were uniformly rejected in naive mice (**Supplementary Fig. 8a, b**) and donor mice (**Fig. 7d, e**).

**Fig. 7.**
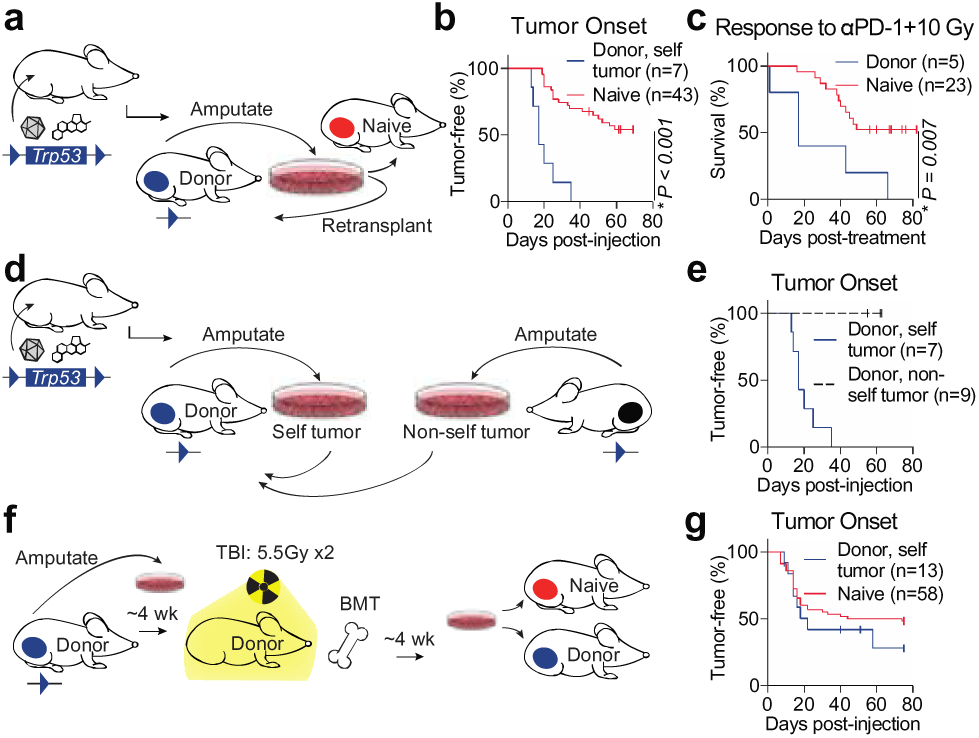
Radiation and αPD-1 fail to overcome immune tolerance. **a**, Transplant tumors generated by injecting cells from a primary tumor into naive mice or donor mice. **b**, Time to tumor onset in donor (blue) or naive (red) mice. **c**, Survival after αPD-1 and 10 Gy RT. **d**, Cell lines from primary tumors in donor (“self”) or other (“non-self”) mice were injected into donor mice. **e**, Time to tumor onset after self (blue) or non-self (black, dashed) cells injected into donor mice. **f**, Cell lines from amputated primary sarcomas were injected into donor or naive mice after TBI and BMT. **g**, Time to tumor onset in donor (blue) or naive (red) mice. Survival curves estimated using Kaplan-Meier method; significance determined by log-rank test and Bonferroni correction.

To test whether this immune tolerance could be overcome with an immune system that did not coevolve with the primary tumor, we again generated primary p53/MCA sarcomas and amputated the tumor-bearing limb when the tumors reached ∼70 mm^3^. We then generated a cell line from each amputated tumor. Approximately 4 weeks later, after the mice had recovered from surgery, they received a lethal dose of total body irradiation (TBI) and rescue by bone marrow transplant (BMT). One month later, donor mice were transplanted with self tumor cells. Naive mice also received a simultaneous injection of the same tumor cells at the time of donor transplant (**Fig. 7f**). Bone marrow transplantation of donor mice restored their ability to reject tumor transplants to the level of naive mice (**Fig. 7f, g**). Taken together, these data demonstrate that primary p53/MCA tumors cause immune tolerance to tumor cells that coevolve with their immune system and cannot be overcome by the immunostimulatory effects of tumor cell injection or by treatment with RT and immunotherapy.

## Discussion

Here, we use high-dimensional profiling to characterize the immune landscapes of autochthonous and transplanted soft tissue sarcomas. We find that primary and transplant tumors derived from the same tumor model have distinc immune landscapes with divergent responses to radiotherapy and anti-PD-1 immunotherapy. To characterize bulk tumor tissue, we performed whole tumor RNA sequencing and whole exome sequencing. We identified distinct transcriptional programs in primary and transplant tumor models. By single-cell sequencing, we reveal transcriptio features of a T cell population that is found in immunotherapy-sensitive transplant tumors. Using mass cytometry, validate that these distinct T cell transcriptional states are reflected at the protein level. By generating autochthono tumors and then performing transplantation experiments, we find that auto-transplantation fails to generate a response to anti-PD-1 immunotherapy. These observations support a model in which coevolution of tumors and th immune system generates an immune cell landscape that favors tumor tolerance, which RT and anti-PD-1 therapy may fail to overcome. Furthermore, we identify features of immune resistance that exist in treatment-resistant prim tumors that co-evolve with the immune system.

Because primary and transplant models of cancer have distinct immune landscapes, they may rely on distinct mechanisms for immunologic clearance. For example, in transplant tumors PD-1 blockade induces a decrease in immunosuppressive M2 macrophages, which is accompanied by an increase in activated T cells. Although similar proinflammatory myeloid cell remodeling occurs in primary tumors after PD-1 blockade, this is not associated with an increase in activated T cells in primary tumors, suggesting that additional mechanisms of immune tolerance are responsible for primary tumor resistance to immunotherapy. Although the responses in transplant tumors suggest synergy with anti-PD-1 therapy and RT, we do not identify synergistic effects to improve survival when mice with primary tumors are treated with combination therapy. Nevertheless, immune checkpoint blockade therapy alone has activity in a subset of cancer patients. Therefore, even if ongoing clinical trials reveal that there is no synergy between radiation and checkpoint blockade in patients, there may be additive effects, which have the potential to improve patient outcomes.

Primary and transplant tumor models may exploit different mechanisms of resistance to immunotherapy and radiation therapy, which has important implications for interpreting preclinical data and translating them to clinical trials. After treatment with anti-PD-1 therapy, transplant but not primary tumors upregulate *Ido1* (**Supplementary Fig. 1g**). Although Ido1 inhibitors enhance the efficacy of anti-PD-1 therapy in transplant mouse models^11^, this combination failed to benefit patients with melanoma^42^. Generating primary tumors and testing their responses to therapy requires a significant investment of time and resources in comparison to studying transplant tumors. However, the effort and resources required to perform these studies in primary tumor models are much less compared to those required for clinical trials, which also have risks for human subjects. Results of cancer therapies in transplant tumor models often fail to predict efficacy for cancer patients^43^. The finding that many cancer therapeutics fail to demonstrate efficacy in clinical trials suggests that many preclinical models do not fully recapitulate the complex nature and diversity of human cancers, which include their interactions with host immune responses, intratumoral heterogeneity, and the diverse cell types present within the tumor microenvironment. Our findings suggest that transplant tumor models may recapitulate the immune microenvironment of highly inflamed tumor subtypes that are likely to respond to immunotherapy, but do not resemble the majority of human cancers, which are resistant to immunotherapy. Performing complementary studies in transplant and autochthonous mouse models may thus increase the success of translation of immunotherapies from preclinical studies to patients.

Over three decades ago, North and Bursuker demonstrated that transplanting tumor cell lines generates an anti-tumor immune response^44^. More recently, Crittenden and colleagues showed that injection of cancer cell lines induces “pre-existing immunity” that is essential for transplant tumor cure by RT and immune checkpoint inhibition^45^. By creating a single cell atlas of immune cells from transplant and primary sarcomas from the same mouse model, we show that tumor cell line injection into syngeneic mice generates a distinct transcriptional state in T cells, characterized by type I and II interferon response programs. Because this T cell response may not occur in the majority of primary tumors, which develop gradually under immunosurveillance, results in transplant models may overestimate human tumor response rates to immunotherapy and RT. Furthermore, upregulation of Wnt/β-Catenin and Myc target gene signaling, which occurs specifically in CD8+ T cells infiltrating primary tumors, may represent a promising therapeutic target in primary tumors that coevolve with the immune system. Identifying this distinct transcriptional program in T cells in primary tumors opens new therapeutic possibilities to enhance the efficacy of immunotherapy and RT for cancer.

## METHODS

### Mouse strains

All animal studies were performed in accordance with protocols approved by the Duke University Institutional Animal Care and Use Committee (IACUC) and adhere to the NIH Guide for the Care and Use of Laboratory Animals. The *Trp53^fl/fl^* allele used in this study has been described previously^46^. *Trp53^fl/fl^* and wild type mice were maintained on a pure 129/SvJae genetic background and bred at Duke University. Nude mice were purchased from Taconic. To minimize the effects of genetic background, age-matched littermate controls were used for every experiment so that potential genetic modifiers would be randomly distributed between experimental and control groups.

### Sarcoma induction and treatment

Primary p53/MCA sarcomas were generated in mice between 6-10 weeks old by intramuscular injection of adenovirus expressing Cre recombinase (Adeno-Cre; University of Iowa Viral Vector Core) into *Trp53^fl/fl^* mice. Twenty-five µL of adenovirus was mixed with 600 µL DMEM (Gibco) and 3 µL 2M CaCl_2_, then incubated for 15 minutes at room temperature prior to injection. Fifty µL of the prepared mixture was injected into the gastrocnemius muscle of the mice, followed by injection of 300 µg MCA (Sigma) resuspended in sesame oil (Sigma) at 6 µg/µL. Transplant p53/MCA sarcomas were generated by injecting 50,000 cells resuspended in 100 µL of a 1:1 mixture of DMEM (Gibco) and matrigel (Corning) into the gastrocnemius muscle.

When tumors reached 70-150 mm^3^ (Day 0, D0), mice were randomized to treatment groups, then tumors were monitored three times weekly by caliper measurements in two dimensions. Antibodies were administered starting on D0 (prior to radiation treatment) by intraperitoneal injection of 200 µL per dose at 1 mg/mL diluted in PBS. Anti-PD-1 (MSD muDX400 in all figures except **Supplementary Fig. 1b**, with BioXCell, BE0146) and anti-CTLA-4 (BioXCell, BE0164) or isotype control (MSD IgG1 control for muDX400, BioXCell, BE0086 for CTLA-4) treatments were administered on days 0, 3, and 6. Anti-CD4 (BioXCell, BE0003-1) and anti-CD8 (BioXCell, BE0061) or isotype control (BioXCell, BE0090) antibodies were injected on day 0, 3, 6, followed by weekly injections for the rest of the experiment. Mice were considered “cured” and censored on Kaplan Meier analysis if tumors were undetectable by caliper measurement for at least 60 days.

Sarcoma irradiations were performed using the Precision Xrad 225Cx small animal image-guided irradiator^47^. The irradiation field was centered on the target via fluoroscopy with 40 kilovolt peak (kVp), 2.5 mA x-rays using a 2 mm aluminum filter and a 0.3 mm copper filter. Sarcomas were irradiated with parallel-opposed anterior and posterior fields with an average dose rate of 300 cGy/min prescribed to midplane with 225 kVp, 13 mA x-rays using a 0.3-mm copper filter and a collimator with a 40 × 40 mm^2^ radiation field at treatment isocenter. Mice were euthanized with CO_2_ if moribund or when tumor volumes reached more than 13 mm in any dimension, in accordance with IACUC guidelines at Duke University.

### Immunohistochemistry

Sarcomas were harvested after euthanasia, fixed in 4% PFA overnight, and preserved in 70% ethanol until paraffin embedding. Formalin-fixed paraffin-embedded tissues were sectioned at 4 µm onto positively-charged slides and baked for 1 hour at 60°C. The slides were then dewaxed and stained on a Leica BOND Rx autostainer (Leica, Buffalo Grove, IL) using Leica Bond reagents for dewaxing (Dewax Solution), antigen retrieval (Epitope Retrieval Solution 2), and rinsing after each step (Bond Wash Solution). Antigen retrieval was performed for 20 minutes at 100°C and all other steps at ambient temperature. Endogenous peroxidase was blocked with 3% hydrogen peroxide for 5 minutes, followed by protein blocking with TCT buffer (0.05M Tris, 0.15M NaCl, 0.25% Casein, 0.1% Tween 20, pH 7.6 +/- 0.1) for 10 minutes. Primary antibody (CD8a, Invitrogen) was applied for 60 minutes followed by secondary antibody (ImmPress HRP Secondary, Vector Labs) application for 12 minutes. Staining was completed with BOND Polymer Refine Detection kit (Leica) including DAB chromogen and hematoxylin nuclear stain. Full-slide images were acquired on the Aperio AT (Leica), and cellular analysis was performed using HALO image analysis software (Indica Labs, Corrales, NM).

### DNA and RNA sequencing

#### RNA sequencing

Tumor specimens and matched liver control were harvested and stored in RNALater (Ambion) at -80°C until all samples were collected. DNA and RNA extractions from each sample were performed using AllPrep DNA/RNA Mini Kit (Qiagen). Extracted total RNA quality and concentration was assessed on a 2100 Bioanalyzer (Agilent Technologies) and Qubit 2.0 (ThermoFisher Scientific), respectively. Only extracts with RNA Integrity Number > 7 were processed for sequencing. RNA-seq libraries were prepared using the KAPA Stranded mRNA-Seq Kit (Roche) following the manufacturer’s protocol. mRNA transcripts were first captured using magnetic oligo-dT beads, fragmented using heat and magnesium, and reverse transcribed using random priming. During the second strand synthesis, the cDNA:RNA hybrid was converted into double-stranded cDNA (dscDNA) and dUTP was incorporated into the second cDNA strand to mark it. second strand. Illumina sequencing adapters were then ligated to the dscDNA fragments and amplified to produce the final RNA-seq library. The strand marked with dUTP was not amplified, allowing strand-specific sequencing. Libraries were indexed using a dual indexing approach so that multiple libraries could be pooled and sequenced on the same sequencing Illumina sequencing flow cell. RNA samples were pooled with whole exome sequencing (WES) libraries for sequencing (see WES section below for additional details). For the RNA samples, sequencing generated an average of approximately 95 million reads per tumor sample.

#### Whole exome sequencing

DNA extraction was followed by RNAse treatment (Qiagen). Genomic DNA samples were quantified using fluorometric quantitation on the Qubit 2.0 (ThermoFisher Scientific). For each sample, 200 ng DNA was sheared using focused-ultrasonicators (Covaris) to generate DNA fragments of about 300 bp in length. Sequencing libraries were then prepared using the Agilent SureSelect XT Mouse All Exon kit (#S0276129). During adapter ligation, unique indexes were added to each sample. Resulting libraries were cleaned using Solid Phase Reversible Immobilization beads and quantified on the Qubit 2.0, and size distribution was checked on an Agilent Bioanalyzer. Libraries were subsequently enriched individually by hybridization of the prepared gDNA libraries with mouse all exome target-specific probes provided with the SureSelect XT Mouse All Exon kit, (target size 49.6 Mb). After hybridization, the targeted molecules were captured on streptavidin beads. Once enriched, the captured libraries were pooled with the RNA libraries and sequenced on an Illumina Novaseq 6000 S4 flow cell at 151 bp paired-end. Base-calling was done on the instrument using RTA v3.3.3. For WES samples, sequencing generated an average of approximately 150 million reads per tumor sample, with tumor coverage at an average depth of 625X, and an average of approximately 38 million reads per liver sample, with liver coverage at an average depth of 157X. Once generated, sequence data were demultiplexed and Fastq files were generated using Bcl2Fastq2 conversion software provided by Illumina (v2.20.0.422).

### RNA sequencing analysis for differential expression

#### Preprocessing

The quality of the sequencing reads was first assessed using FastQC (v0.11.5)^48^ and MultiQC^49^. Low quality reads and adapters were detected and removed with Trimmomatic (v0.36)^50^. The quality of the reads was assessed again before downstream analyses and qualified reads were then aligned to UCSC mm10 mouse genome from the iGenome project using STAR (v2.5.4b)^51^ and mapped to the mouse transcriptome annotated by GENCODE (Release M17)^52^. Summaries for alignment and mapping performance are provided in **Supplementary Table 4**. Raw gene counts were quantified using HTSeq^53^ implemented in the STAR pipeline.

#### Gene Differential Expression Analysis

Normalization of gene counts and differential expression analysis were performed based on modeling the raw counts within the framework of a negative binomial model using the R package DESeq2 (v1.20.0)^54^. Pathway analyses, based on Gene Ontology (GO) terms were conducted using the gage package (v2.34.0)^55^. The Benjamini-Hochberg method^56^ was used to adjust p values for multiple testing within the false-discovery framework.

#### Gene Signature Analysis

A gene signature that responds to interferons (ISG signature) was obtained from Liu *et al*^21^. Human genes were converted to mouse equivalent genes using the Ensembl database (Release 97)^57^, which resulted in a final set of 44 genes(*Adar*, *Bst2*, *Casp1*, *Cmpk2*, *Cxcl10*, *Ddx60*, *Dhx58*, *Eif2ak2*, *Epsti1*, *Herc6*, *Ifi35*, *Ifih1*, *Ifit2*, *Ifit3*, *Ifit3b*, *Irf7*, *Isg15*, *Isg20*, *Mx2*, *Mx1*, *Nmi*, *Oasl1*, *Ogfr*, *Parp12*, *Parp14*, *Pnpt1*, *Psme2*, *Psme2b*, *Rsad2*, *Rtp4*, *Samd9l*, *Sp110*, *Stat2*, *Tdrd7*, *Trafd1*, *Trim14*, *Trim21*, *Trim25*, *Ube2l6* and *Usp18*). The same modified Z-scores^21^ were applied to the normalized expression data to summarize the expression level of the ISG signature. One-way ANOVA model and Tukey’s multiple comparison test were used to compare the signature scores between all pairs of the treatment groups.

All statistical analyses related to differential expression were performed in the R statistical environment along with its extension packages from the comprehensive R archive network (CRAN; https://cran.r-project.org/) and the Bioconductor project^58^.

### RNA-sequencing analysis for neoantigen expression analysis

Alignment of RNA sequencing data was performed in Omicsoft Array Studio (v10.0.1.118). Briefly, cleaned reads were aligned to the mouse B38 genome reference by using the Omicsoft Sequence Aligner (OSA)^59^, with a maximum of two allowed mismatches. Gene level counts were determined by the OSA algorithm as implemented in Omicsoft Array Studio and using Ensembl.R86 gene models. Approximately 88% of reads across all samples mapped to the mouse genome (corresponding to approximately 84 million reads).

### Whole exome sequencing analysis

#### Somatic mutation calling

The WES reads were aligned to the mouse reference genome mm10 using the BWA-MEM algorithm (v0.7.12)^60^. The reference genome was obtained from the UCSC FTP site (ftp://hgdownload.soe.ucsc.edu/goldenPath/mm10/). The original Agilent bait designed BED file, which had been built on GRCm37 (mm9), was lifted over to GRCm38 (mm10) to match the other reference files.

The aligned bam files were post-processed by following the recommended pipeline of Genome Analysis Toolkit (GATK, version 3.8)^61^ to generate the analysis-ready BAM files for variant calling. Briefly, first, Picard (v1.114; http://broadinstitute.github.io/picard/faq.html) MarkDuplicates module was used for identifying PCR duplication. Afterwards, the reads were locally realigned around insertion or deletion (indels) by module RealignerTargetCreator/IndelRealigner of GATK. Finally, module BaseRecalibrator of GATK was performed to recalibrate quality scores.

Somatic mutations were detected using GATK3.7 MuTect2^62^ with default parameters by inputting the analysis-ready BAM files of tumor and matched normal tissue (liver) control for each animal (parameter “--dbsnp” was assigned with mouse Single Nucleotide Polymorphism Database (dbSNP) (v150)^63^. Variants called by MuTect2 that were present in the dbSNP were removed. Variants with a mutant allele depth <4 or total read depth <15 were excluded. Variants were annotated with their most deleterious effects on Ensembl transcripts with Ensembl VEP (Variant Effect Predictor, Version 88)^64^ on GRCm38.

#### Neoantigen prediction

Mutant 8-11mer peptides that could arise from the identified non-silent mutations present in each tumor were identified. If the variant gave rise to a single amino acid change, the mutant peptide was scanned with a sliding window of 8–11 amino acids around the variant to generate all possible 8, 9, 10 and 11mers. If the variant created large novel stretches of amino acids that were not present in the reference genome (e.g. stop losses, or frameshifts), all possible peptides of 8, 9, 10 and 11mers were extracted from the large novel peptide. The binding ability between all the mutant peptides and mouse H2-K^b^/D^b^ were predicted by netMHC (4.0)^65^ with default parameters. For each specific variant, the associated peptides were considered to be neoantigens if they met the following criteria: the half-maximal inhibitory concentration binding affinity scores (affinity score) of mutant peptides < 500 nM and reference peptide affinity score > 500 nM. The number of RNA-Seq reads covering each of the predicted variants was extracted from the samtools^66^ (v1.6) mpileup results (with RNA-Seq bam files processed by Omicsoft (please see section “RNA-sequencing analysis for neoantigen expression” above). Neoantigens were considered expressed if both the mutant and the reference alleles were found in at least five RNA-seq reads.

#### Genomic immune editing and transcriptional neoantigen depletion

DNA immunoediting was identified by calculating the observed-to-expected ratio of neoantigens^41, 67^. Neoantigens derived from indels were excluded, and only neoantigens from missense mutations were considered observed. The neoantigens derived from silent mutations were considered expected neoantigens. For silent mutations, reference peptide affinity score was not included in the neoantigen criteria. The observed-to-expected ratio of neoantigens was calculated as the number of neoantigens per missense mutation divided by the number of neoantigens per silent mutation. Transcriptional neoantigen depletion was calculated by dividing the number of expressed neoantigens (as defined above) by the total number of neoantigens.

### Single-cell RNA sequencing

#### Tumor harvest and dissociation

Tumors were dissected from mice, minced, and digested using the Miltenyi Biotec tumor dissociation kit (mouse, tough tumor dissociation protocol) for 40 minutes at 37°C. Cells were then strained through a 70 µm filter and washed with FACS buffer (HBSS (Gibco) with 5 mM EDTA (Sigma-Aldrich) and 2.5% fetal bovine serum (Gibco)). Red blood cells were lysed using ACK lysis buffer (Lonza), washed again with FACS buffer, and strained through a 40 µm filter. Cells were then washed and stained for cell sorting.

#### Fluorescence-activated cell sorting

For sorting of CD45+ cells for single-cell RNA sequencing, single-cell suspensions of tumors were blocked with purified rat anti-mouse CD16/CD32 (BD Pharmingen, dilution 1:100) for 10 minutes at room temperature then stained with Live/Dead dye (Zombie Aqua, Biolegend) and anti-mouse CD45 (BV605 or APC-Cy7, Biolegend) for 25 minutes on ice. Live CD45+ cells were isolated for scRNA-seq using an Astrios (Beckman Coulter) sorter and resuspended in PBS with 0.04% BSA at a concentration of 1000 cells/µL for single-cell RNA sequencing.

#### Library preparation and sequencing

scRNA-seq was performed as previously described^68^. Single-cell suspensions from sorted live CD45+ cells were loaded on a GemCode Single Cell instrument (10x Genomics) to generate single cell beads in emulsion and scRNA-seq libraries were prepared using the Chromium Single Cell 3’ Reagent Kits (v2), including Single Cell 3′ Library & Gel Bead Kit v2 (120237), Single Cell 3 Chip Kit v2 (PN-120236) and i7 Multiplex Kit (120262) (10x Genomics) following the Single Cell 3 Reagent Kits (v2) User Guide. Single cell barcoded cDNA libraries were quantified by quantitative PCR (Kappa Biosystems) and sequenced on an Illumina NextSeq 500 (Illumina). Read lengths were 26 bp for read 1, 8 bp for i7 index, and 98 bp for read 2. Cells were sequenced to greater than 50,000 reads per cell as recommended by manufacturer.

#### Analysis of scRNA-seq data

The CellRanger Single Cell Software Suite (v2.1.1) was used to perform sample de-multiplexing, barcode processing and single cell 3’ gene counting. Reads were aligned to the mouse (mm10) reference genome. Cells that had less than 200 expressed genes, more than 7,000 expressed genes, or greater than 0.05% of mitochondrial genes were excluded from analysis. Graph-based cell clustering, dimensionality reduction, and data visualization were analyzed by the Seurat R package^69^ (v2.4). The number of cell clusters is determined using graph-based clustering in Seurat that embeds cells in a K-nearest neighbor graph based on the Euclidean distance in PCA space and Jaccard similarity to iteratively group cells together with similar gene expression patterns. We determined the number of statistically significant principal components to input into the graph-based clustering algorithm using jackStraw^70^, but selected cluster resolution according to Seurat^71^ recommendations (https://satijalab.org/seurat/v3.0/pbmc3k_tutorial.html) based on the size of the dataset . We used tSNE visualization to confirm appropriate clustering. Differentially expressed transcripts were determined in the Seurat package utilizing a likelihood-ratio test for single-cell gene expression^72^. Graphics were generated using Seurat, ggplot2 R packages, and Graphpad Prism (v8). To identify subclusters within the lymphoid and myeloid cell compartments we reanalyzed selected cells within lymphoid and myeloid clusters using the Seurat pipeline described above.

#### Gene set variation analysis (GSVA)

To identify functionally enriched pathways between tumor types or between treated and untreated tumor in CD8+ T Cells, we used a modified version of previously described methods^73^. First, to assign pathway activity estimates to individual cells, we applied GSVA^74^ to calculate enrichment scores in each cell for the 50 hallmark pathways described in the molecular signature database^75^ (MSigDB v6.0), as implemented in the GSVA R package (v1.32.0). We then tested each pair of conditions for a difference in the GSVA enrichment scores of each hallmark pathway. We used a simple linear model and t statistics implemented in the limma^76^ R package (v3.38.3) that uses an empirical Bayes shrinkage method. The Benjamini-Hochberg method^56^ was used to adjust p values for multiple hypothesis testing within the false-discovery framework.

#### Cell type annotation

The R package *SingleR* v1.0.1^77^ was used to annotate cell types based on correlation profiles with bulk RNA-seq from the Immunological Genome Project (ImmGen) database^78^. Cell type annotation was performed for individual cells of the whole dataset, as well as the lymphocyte and myeloid cell subsets of the dataset.

### Mass Cytometry

#### Staining

Antibody clones and sources are listed in **Supplementary Table 5**. For custom-conjugated antibodies, 100 µg of antibody was coupled to Maxpar X8 metal-labeled polymer according to the manufacturer’s protocol (Fluidigm). After conjugation, the metal-labeled antibodies were diluted in Antibody Stabilizer PBS (Candor Bioscience) for long-term storage. After tumor dissociation and RBC lysis as described above, three million cells per sample were transferred to 5 mL round-bottom tubes (Corning). Cells were incubated with 300 µL of Cell-ID Cisplatin-195Pt (Fluidigm) diluted 1:4000 in Maxpar PBS (Fluidigm) for 5 minutes at room temperature, then washed twice with Maxpar Cell Staining Buffer (CSB) (Fluidigm). Samples were incubated with 50 µL FcR Blocking Reagent (Biolegend, 1:100 dilution) for 10 min at room temperature, then 50 µL extracellular antibody cocktail was added and incubated for 30 minutes total at room temperature. Cells were washed twice with CSB, then fixed and permeabilized with eBioscience Foxp3/Transcription Factor Fixation/Permeabilization Buffer for 1 hour at room temperature, followed by two washes with permeabilization buffer (eBioscience). Fifty µL intracellular antibody cocktail in permeabilization buffer was added and incubated for 30 minutes total at room temperature, followed by two washes with permeabilization buffer. Cells were fixed in 1.6% methanol-free PFA (Thermofisher) diluted with Maxpar PBS (Fluidigm) for 1 hour at 4°C. Each sample was barcoded with a unique combination of palladium metal barcodes (Fluidigm) for 30 minutes. Samples were washed twice, combined, and incubated at least overnight in Maxpar Fix and Perm Buffer (Fluidigm) with 62.5 nM Cell-ID Intercalator (Fluidigm) containing ^191^Ir and ^193^Ir. Before collection, cells were washed once with CSB, once with Cell Acquisition Solution (CAS) (Fluidigm), then filtered and diluted in CAS containing 10% EQ Calibration Beads (Fluidigm) at 0.5 million cells per mL before acquisition on a mass cytometer (Helios).

#### CyTOF data analysis

Mass cytometry data were normalized, concatenated, and debarcoded using Fluidigm CyTOF software (v7.0). Individual samples were gated in Cytobank^79^ to exclude beads, debris, dead cells, and doublets for further analysis. For each experimental group (time point and treatment), cells from 2-4 tumors per group were manually gated to identify specific populations. To generate viSNE plots, all live CD45+ cells from each group were concatenated into a single file using CATALYST^80^. Within Cytobank, viSNE^81^ was applied to 40,000 events per group, followed by FlowSOM^82^ analysis.

### Tumor Transplantation Experiments

For amputation and tumor transplantation experiments, primary tumors were generated using Adeno-Cre and MCA as described above. After hind limb amputation, tumors were dissected from the limb and dissociated by shaking for 45 minutes at 37°C in collagenase Type IV (Gibco), dispase (Gibco), and trypsin (Gibco). Cell suspension was then strained through a 40 µm filter, washed in PBS, and plated for culture. Cell lines were maintained *in vitro* for 10 passages before transplanting into naive or donor mice. For each donor mouse that was transplanted with a “self” cell line, approximately 5 matched naive mice also received an injection of the same cell line. For both donor and naive mice, transplant sarcomas were generated by injecting 50,000 cells resuspended in 100 µL of a 1:1 mixture of DMEM (Gibco) and matrigel (Corning) into the gastrocnemius muscle.

#### Bone marrow transplant

Mice receiving bone marrow transplant were treated with 2 fractions of 5.5 Gy total body irradiation (TBI) delivered 18 hours apart. TBI was delivered using the Precision Xrad 225Cx small animal image-guided irradiator with no collimator and parallel-opposed anterior and posterior fields. Bone marrow transplant was performed within 3 hours of the second TBI dose. Whole bone marrow cells were isolated from femurs and tibias of healthy mice on a 129/SvJae genetic background by washing the bone marrow space with PBS. RBCs were lysed using ACK lysing buffer (Lonza). Bone marrow cells were counted with an automated cell counter (Cellometer Auto 2000, Nexcelom Bioscience) using AO/PI stain (Nexcelom Bioscience). Three million whole bone marrow cells were resuspended in 50 µL PBS and injected retro-orbitally into recipient mice.

### Statistics and study design

Experiments were designed such that littermate controls were used for all experiments. Statistical tests performed are indicated in figure legends. The experiments were randomized, and investigators were blinded to treatment during measurement and data collection. No statistical methods were used to predetermine sample size. Measurements were taken from distinct samples; the same sample was not measured repeatedly.

### Reagent information

See **Supplementary Table 5** for reagent catalogue numbers, antibody clones, and dilutions.

### Code availability

Computer codes used to generate results in this manuscript can be found at gitlab.oit.duke.edu (Project ID 13688). The analysis for neoantigen calling was performed with a proprietary pipeline and we are unable to publicly release this code. We have released all raw data and the implementation details in the Methods and Supplementary Information to allow for independent replication.

## SUPPLEMENTARY INFORMATION

Additional information can be found in the 8 Supplementary Figures and 5 Supplementary Tables.

## Acknowledgments

We thank Anton Berns (Netherlands Cancer Institute) for providing the *Trp53^fl/fl^* mice and Tyler Jacks (MIT) for providing the 129/SvJae mice. Mass cytometry analysis was performed at the University of North Carolina Mass Cytometry Core Lab, which is supported by the University Cancer Research Fund (UCRF) and the UNC Cancer Center Core Support Grant (P30CA016086). We thank David Corcoran (Duke Genomic Analysis and Bioinformatics Shared Resource) for providing valuable bioinformatics support and feedback. We thank MSD for providing muDX400 and isotype control antibodies. We thank Charles Drake for providing critical feedback on this manuscript. This work was supported by awards F30CA221268 (AJW), ASCO Young Investigator Award (YMM), U24 CA220245 (YMM), RSNA Research Resident/Fellow Grant (YMM), Sarcoma Alliance for Research through Collaboration (SARC) SPORE 5U54CA168512 (DGK and YMM), T32GM007171 (AJW and JEH), R35CA197616 (DGK), The Leon Levine Foundation (DGK), the Emerson Collective (DGK), the Duke Cancer Center Support Grant (P30CA14236), and P01CA142538 (KO) from the National Cancer Institute. The funders had no role in the design of the study, collection, analysis, and interpretation of the data or writing the manuscript.

## Author contributions

AJW, YMM, and DGK conceived the project, designed experiments, and wrote the manuscript. AJW, YMM, CSH, JEH, ESX, DJC, CLK, and LL bred and genotyped mice. AJW, YMM, CSH, JEH, DJC, CLK, and ESX induced and measured sarcomas and treated mice with antibody. AJW, YMM, and NW performed radiation treatments. AJW and CSH harvested tissue, generated cell lines, and extracted DNA and RNA for RNA-seq and WES. XQ, DZ, and KO performed analysis of all mouse survival data and RNA-seq data and deposited the RNA-seq and WES data to publicly available databases. LC and ESM performed analysis of RNA-seq data and WES data for mutational load and neoantigen analysis. AJW performed and analyzed CyTOF and collected and processed tumor cells for scRNA-seq. TB performed and analyzed scRNA-seq data and uploaded scRNA-seq data to the SRA database. HF performed GSVA analysis on scRNA-seq data. AJW and CSH collected tumors for histology and YM processed tissue for histology. KSS performed immunohistochemistry. AJW, YMM, CSH, and ESX performed surgery on mice. AJW performed bone marrow transplants. All authors reviewed the manuscript.

## Data availability

All sequencing data have been deposited in publicly accessible databases: bulk tumor RNA-seq (NCBI Gene Expression Omnibus (GEO) database, accession number GSE134273); whole exome sequencing (Bioproject database, project number PRJNA556574); and scRNA-seq (SRA, accession number PRJNA556477). Mass cytometry data are available at flowrepository.org (ID FR-FCM-Z28C). All other relevant data are available from the corresponding authors upon reasonable request.

### Competing interests

DGK is a cofounder of XRAD Therapeutics, which is developing radiosensitizers. DGK and YMM are recipients of a Stand Up To Cancer (SU2C) MSD Catalyst Grant studying pembrolizumab and radiation therapy in sarcoma patients. DGK has received research funding from XRAD Therapeutics, Eli Lilly & Co., and served as chair of the Developmental Therapeutics Committee for the Sarcoma Alliance for Research through Collaboration. ESM and LC are employees of Merck Sharp & Dohme Corp, a subsidiary of Merck & Co., Inc., Kenilworth, NJ, USA. The other authors declare no competing financial interests.

**Correspondence and requests for materials** should be addressed to YMM or DGK.

## SUPPLEMENTAL FIGURES

**Supplementary Figure 1.**
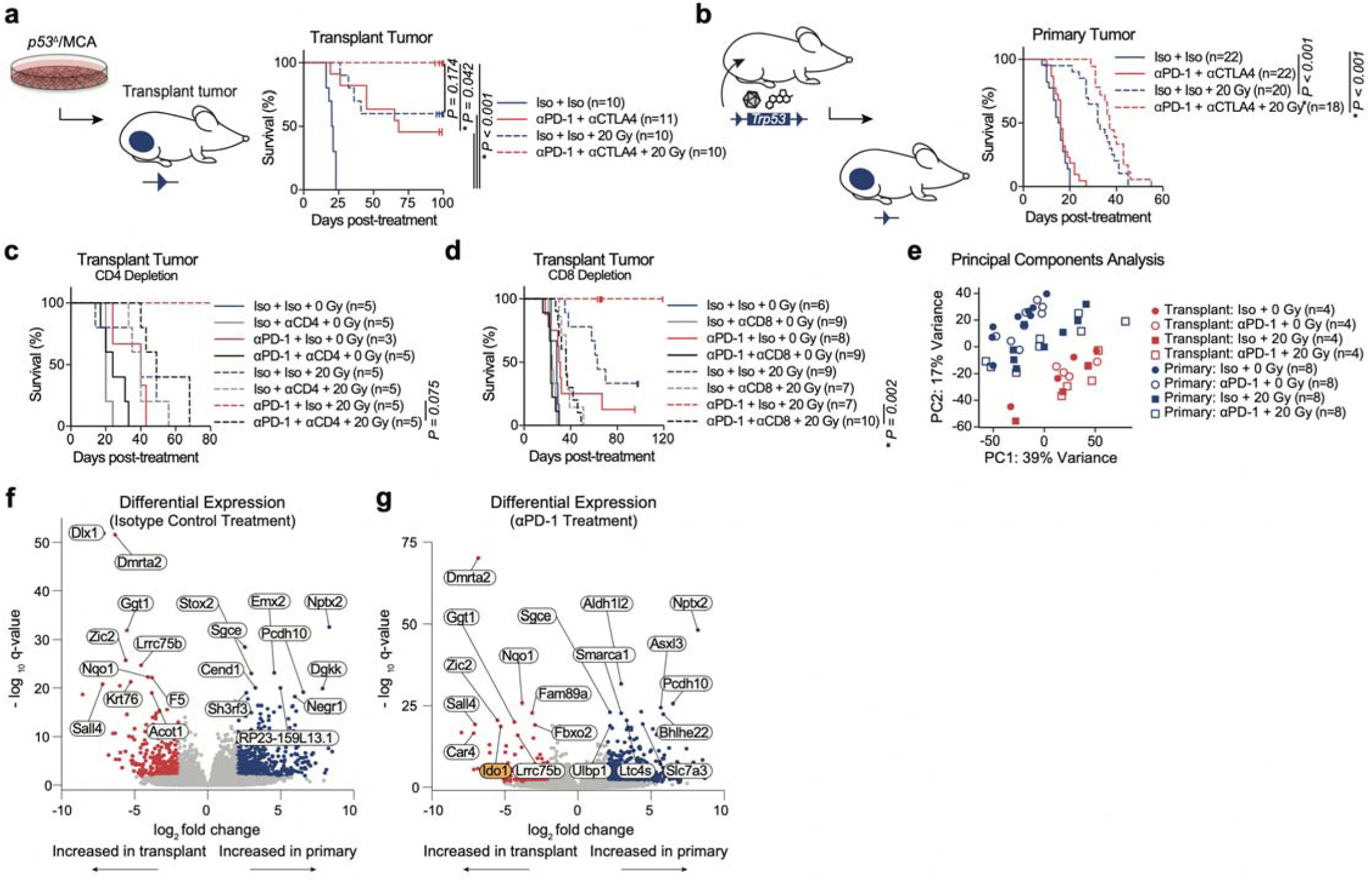
Response of transplant and primary tumors to radiation and immunotherapy. **a**, Mice with transplant sarcomas received either αPD-1 and αCTLA-4 (red) or isotype control (blue) antibodies and 0 (solid) or 20 (dashed) Gy radiation when tumors reached >70 mm^3^. **b**, Mice with primary sarcomas received treatment as in panel a. **c**, Mice with transplant sarcomas received either αPD-1 (red) or isotype control (blue) antibodies, αCD4 (gray) or isotype control (black), and 0 (solid) or 20 (dashed) Gy radiation when tumors reached >70 mm^3^. Surviving mice were censored 164 days after treatment. **d**, Mice with transplant sarcomas received either αPD-1 (red) or isotype control (blue) antibodies, αCD8 (gray) or isotype control (black), and 0 (solid) or 20 (dashed) Gy radiation when tumors reached >70 mm^3^. **e**, Principal components analysis of bulk tumor RNA sequencing harvested 3 days after treatment. Each symbol represents an individual mouse’s tumor. **f**, Differentially expressed genes in isotype control-treated primary (blue) tumors relative to transplant (red) tumors. **g**, Differentially expressed genes in αPD-1-treated primary (blue) tumors relative to transplant (red) tumors. For **a**-**d**, significance determined by log-rank test and Bonferroni correction.

**Supplementary Figure 2.**
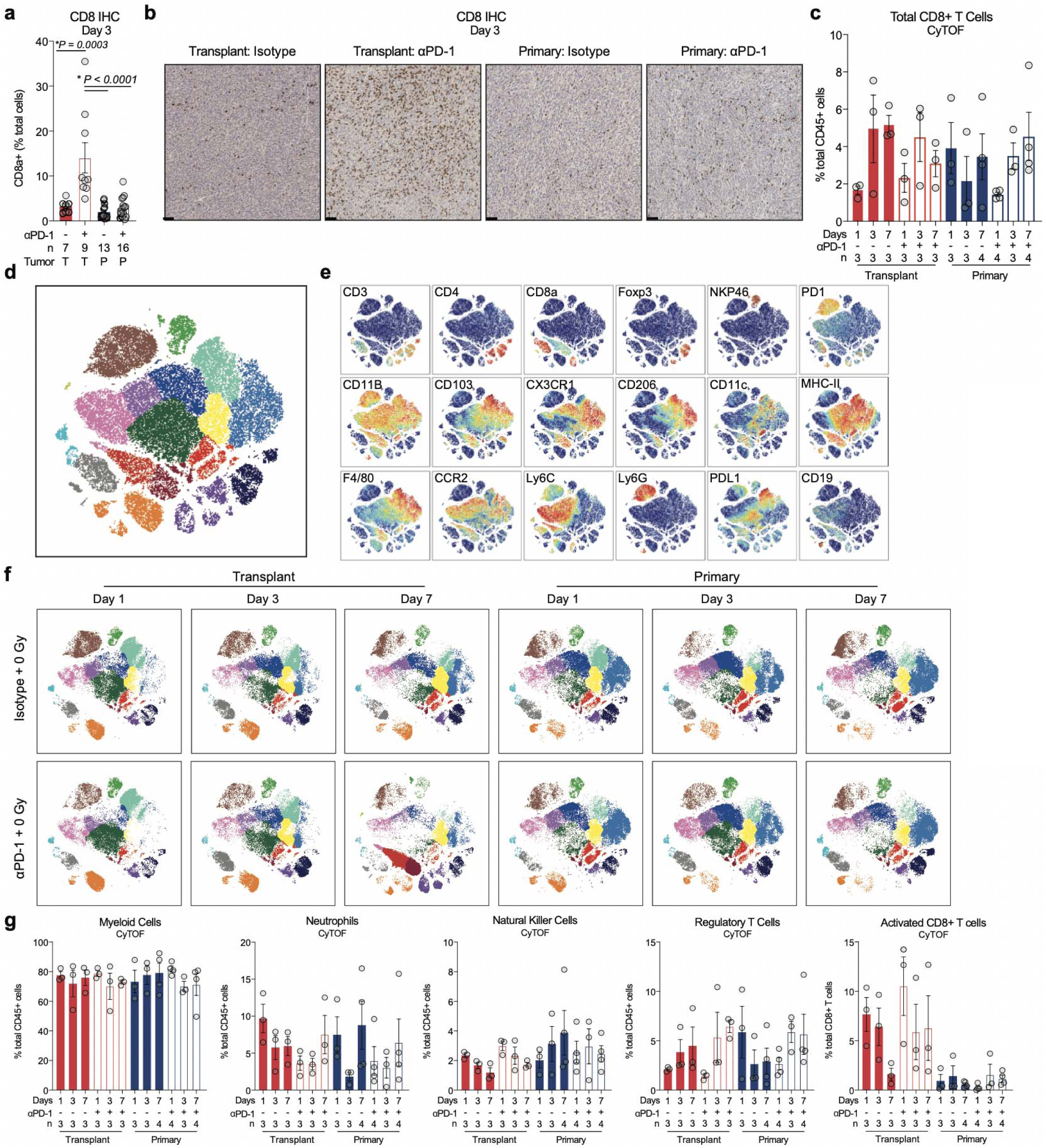
PD-1 blockade increases T Cells in transplant, but not primary sarcomas. **a**, Percent CD8a+ cells by immunohistochemistry 3 days after treatment. T, transplant; P, primary. Significance determined by one-way ANOVA with Tukey’s multiple comparisons test. **b**, Representative histology of CD8a IHC. Scale bars = 100 µm. **c**, Quantification of CD8+ T cell frequency by CyTOF after αPD-1 or isotype control. **d**, CD45+ immune cells from indicated tumor type and treatment groups at 1, 3, and 7 days after treatment, overlaid on viSNE representation and color coded by cluster. **e**, Expression of selected marker proteins. **f**, Distribution of immune cells, separated by indicated tumor type, treatment, and time point. **g**, Bar graphs comparing frequency of indicated cell type (% CD45+) from transplant or primary tumors harvested 1, 3, or 7 αPD-1 (+) or isotype control (-). Data are mean ± SEM.

**Supplementary Figure 3.**
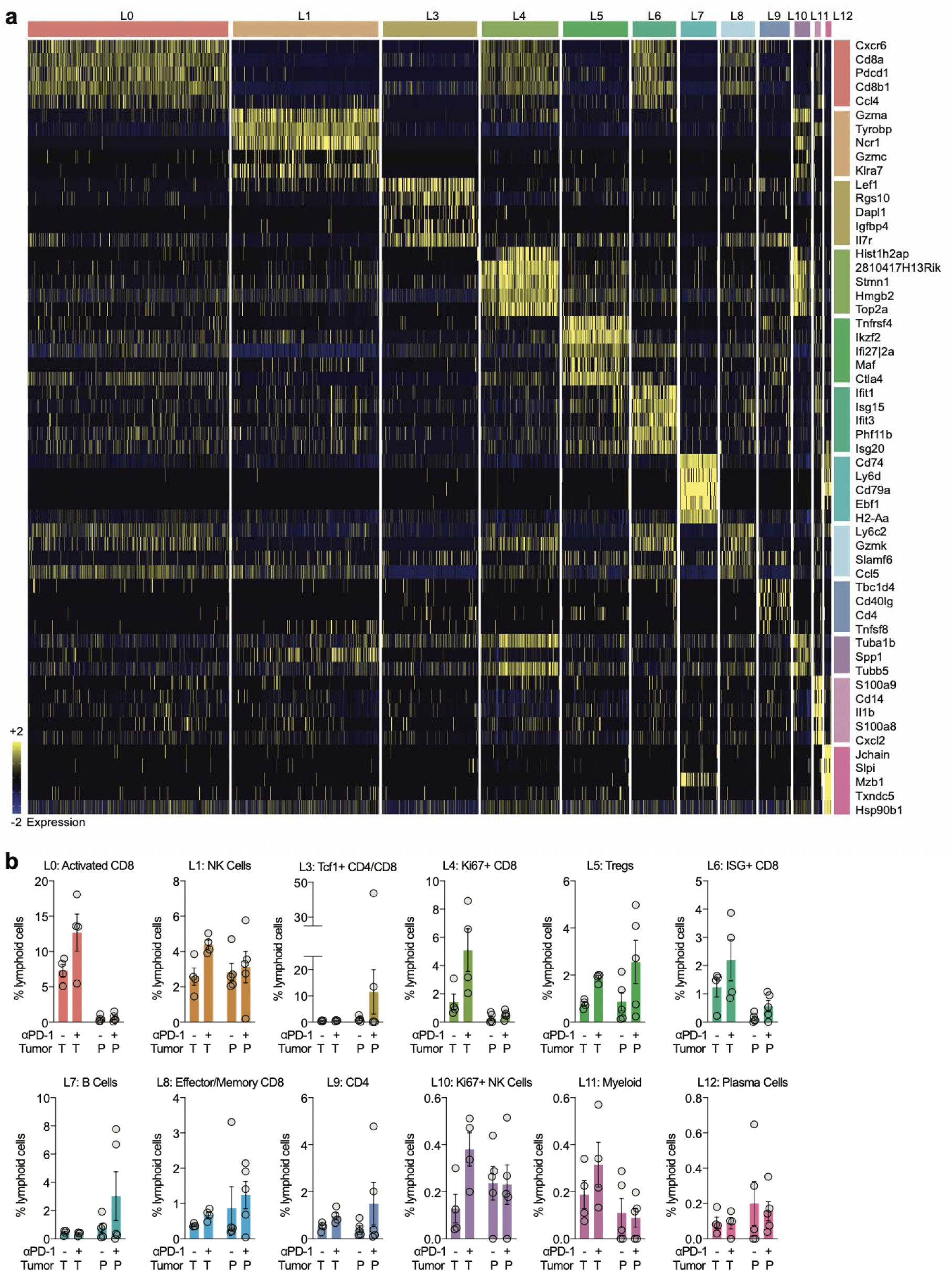
Defining lymphoid populations using scRNA-seq. **a**, Scaled expression values of top three discriminative genes per cluster. **b**, Bar graphs comparing frequency of indicated cell type (% lymphoid cells) from transplant (T) or primary (P) tumors after αPD-1 (+) or isotype control (-). Each symbol represents an individual mouse; n = 4 per group for transplant tumors, n = 5 per group for primary tumors. Data show mean ± SEM.

**Supplementary Figure 4.**
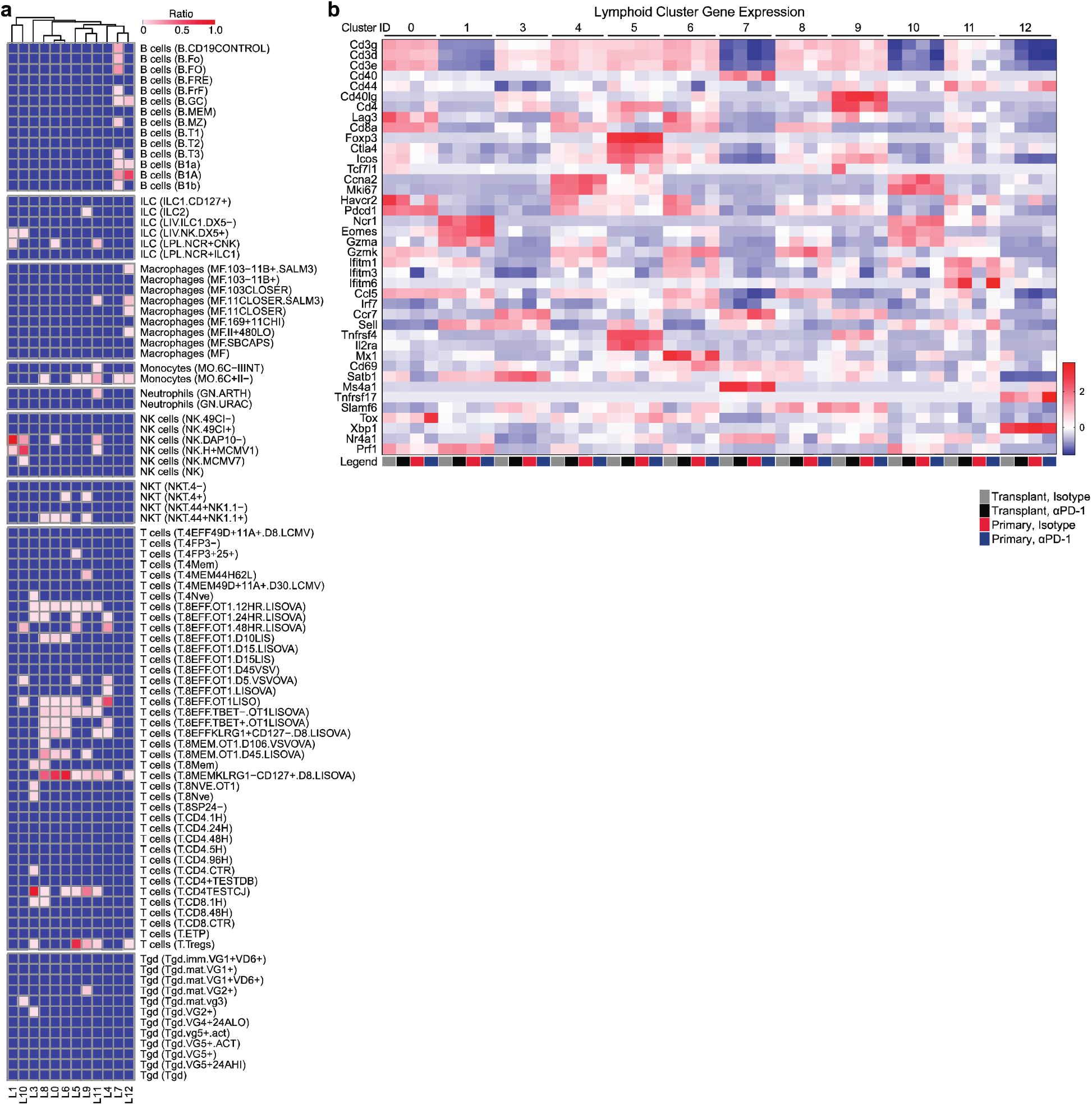
Transcriptional phenotypes of lymphoid populations. **a**, SingleR analysis of lymphoid clusters. The heatmap color scale corresponds to the ratio of cells attributed to that cell type in each cluster. **b**, Heat map comparing expression of genes in indicated clusters and treatment groups.

**Supplementary Figure 5.**
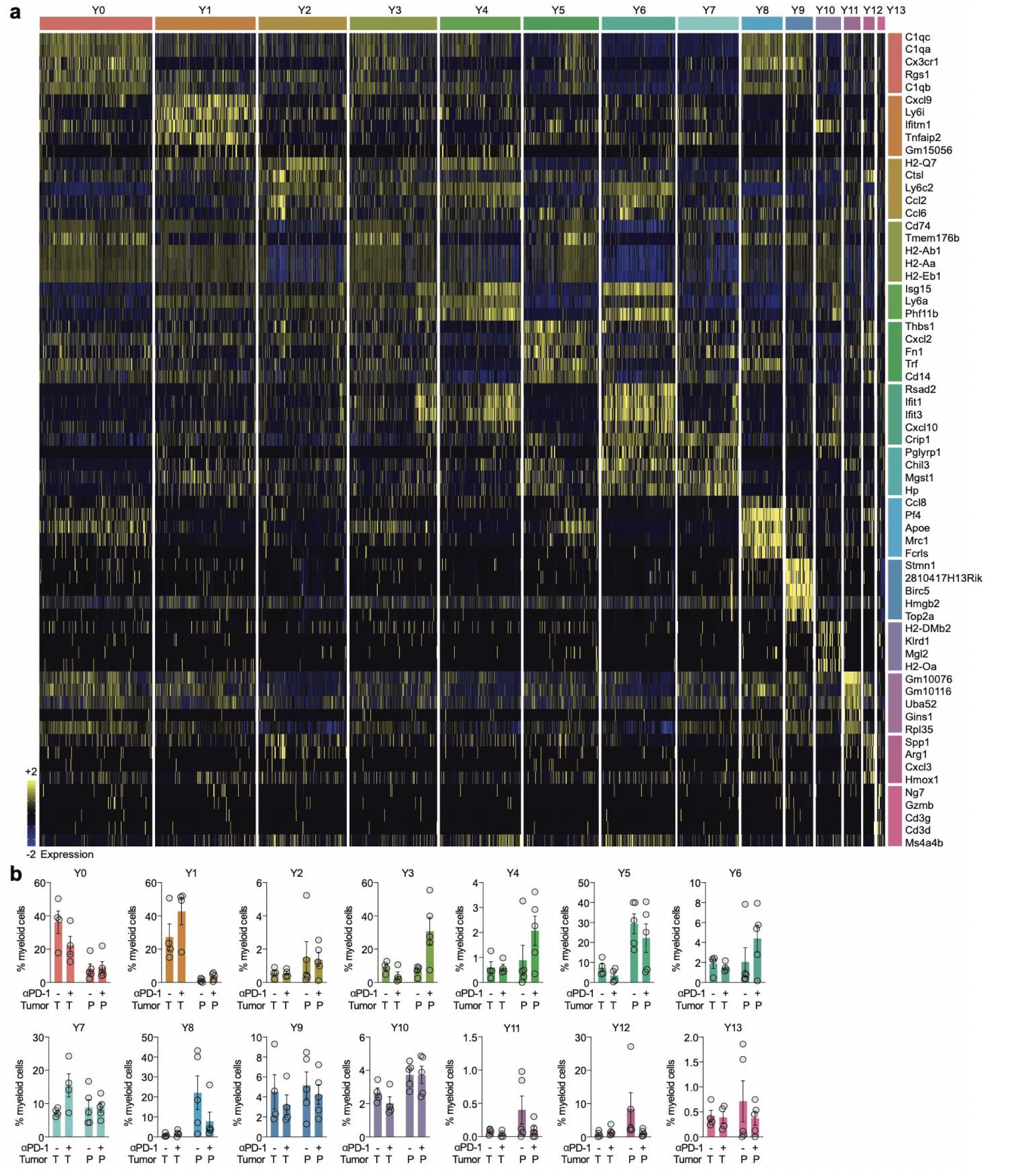
Defining myeloid populations using scRNA-seq. **a**, Scaled expression values of top discriminative genes per cluster from myeloid cell clusters. **b**, Bar graphs comparing frequency of indicated cell type (% myeloid cells) from transplant (T) or primary (P) tumors after αPD-1 (+) or isotype control (-). Each symbol represents an individual mouse; n = 4 per group for transplant tumors, n = 5 per group for primary tumors. Data show mean ± SEM.

**Supplementary Figure 6.**
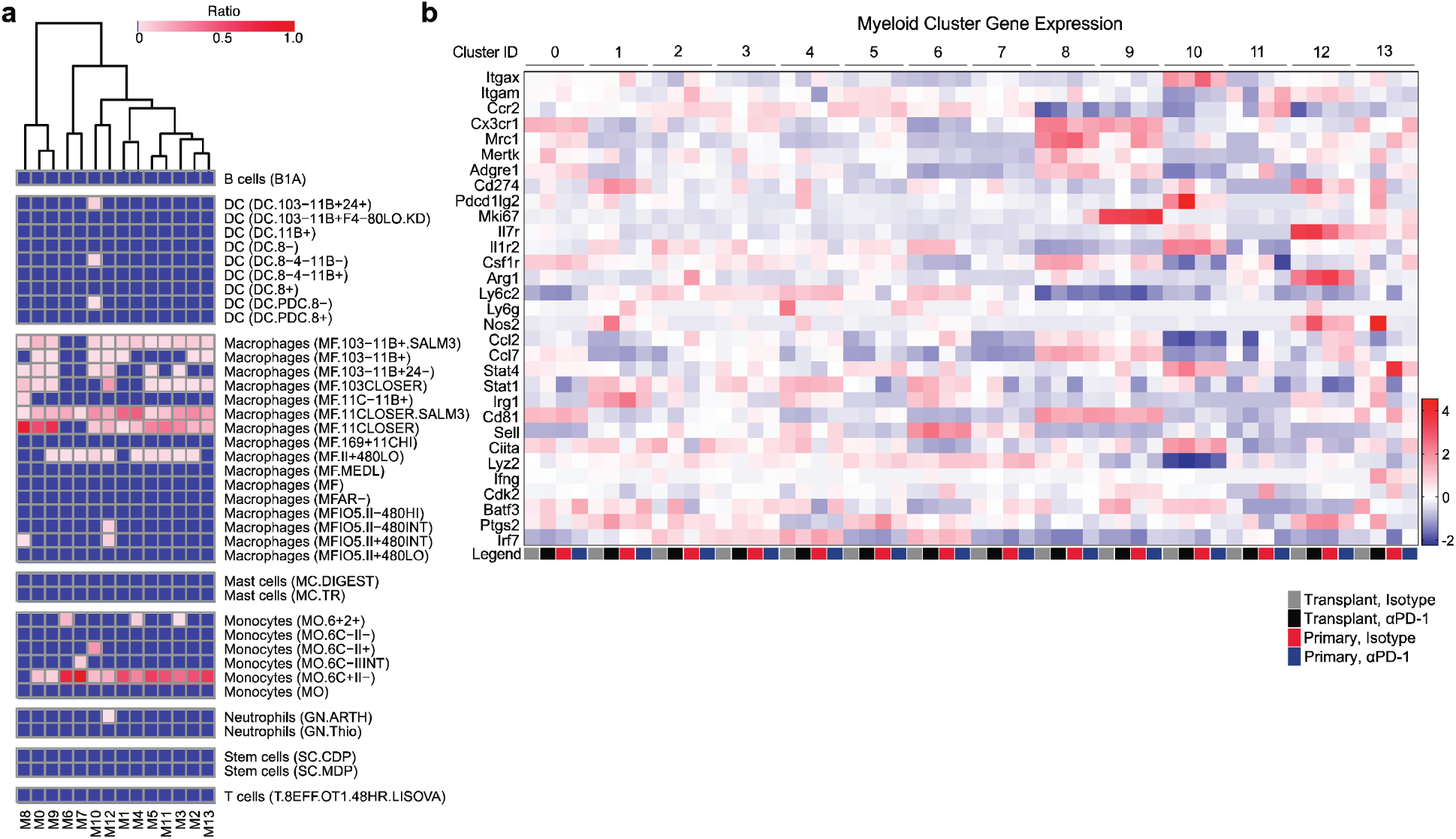
Transcriptional phenotypes of myeloid populations. **a**, SingleR analysis of myeloid clusters. The heatmap color scale corresponds to the ratio of cells attributed to that cell type for each cluster. **b**, Heat map comparing expression of genes in indicated clusters and treatment groups.

**Supplementary Figure 7.**
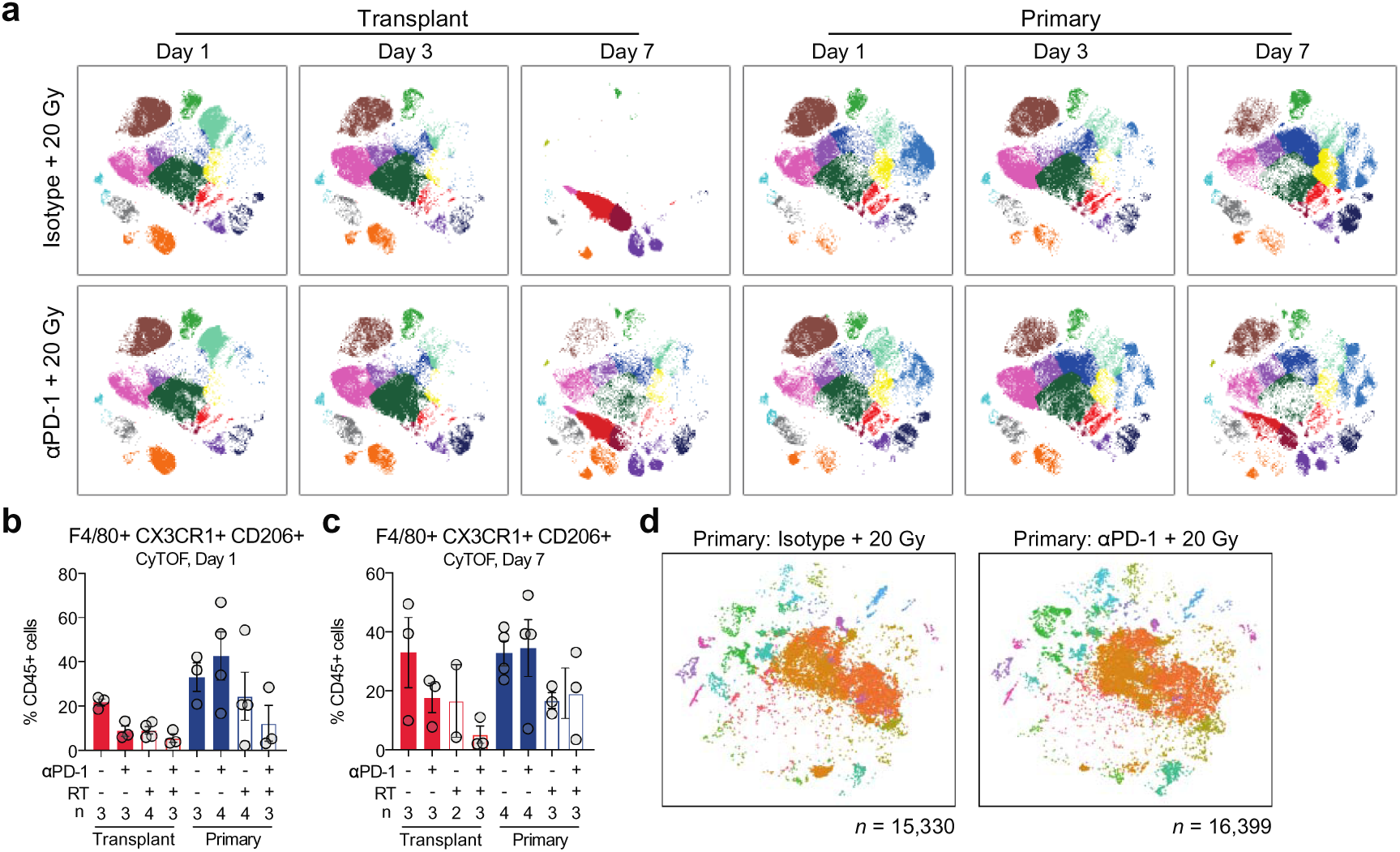
Profiling immune effects of radiation therapy by mass cytometry and scRNA-seq. **a**, CD45+ immune cells from indicated tumor and treatment groups at 1, 3, and 7 days after treatment, overlaid on viSNE representation and color coded by cluster. **b**, Cell frequency by CyTOF one day after treatment. **c**, Cell frequency by CyTOF seven days after treatment. **d**, scRNA-seq of CD45+ immune cells from indicated treatment groups, overlaid on tSNE representation and color coded by cluster.

**Supplementary Figure 8.**
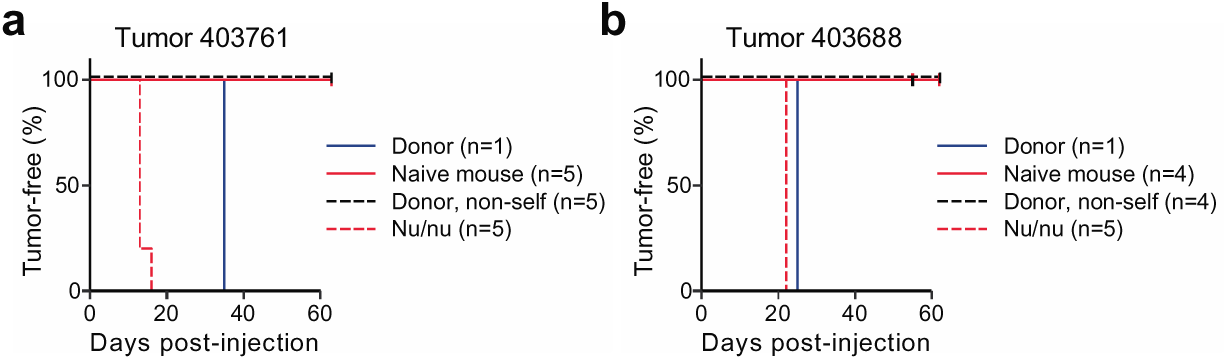
Tumor cell line rejection by immunocompetent mice. **a**, Time from cell line injection to tumor onset in indicated mice for tumor 403761. **b**, Same as **a**, for tumor 403688. Nu/nu = nude mouse.

